# Local angiogenic interplay of Vegfc/d and Vegfa drives brain region-specific development of fenestrated capillaries

**DOI:** 10.1101/2022.12.08.519692

**Authors:** Sweta Parab, Olivia A. Card, Qiyu Chen, Luke D. Buck, Rachael E. Quick, William F. Horrigan, Gil Levkowitz, Benoit Vanhollebeke, Ryota L. Matsuoka

**Affiliations:** Department of Neurosciences, Lerner Research Institute, Cleveland Clinic, Cleveland, USA; Department of Molecular Medicine, Cleveland Clinic Lerner College of Medicine, Case Western Reserve University, Cleveland, USA; Departments of Molecular Cell Biology and Molecular Neuroscience, The Weizmann Institute of Science, Rehovot, Israel; Laboratory of Neurovascular Signaling, Department of Molecular Biology, ULB Neuroscience Institute, Université libre de Bruxelles (ULB), Gosselies, Belgium; Walloon Excellence in Life Sciences and Biotechnology (WELBIO), Wavre, Belgium

**Keywords:** brain angiogenesis, central nervous system, vascular heterogeneity, fenestrated endothelial cells, choroid plexus, pineal gland, pituitary, choriocapillaris, Vegfs, zebrafish

## Abstract

Fenestrated and blood-brain barrier (BBB)-forming endothelial cells constitute major brain capillaries, and this vascular heterogeneity is crucial for region-specific neural function and brain homeostasis. How these capillary types emerge in a brain region-specific manner and subsequently establish intrabrain vascular heterogeneity remains unclear. Here, we show a core angiogenic mechanism critical for fenestrated brain capillary development via a comparative analysis of the zebrafish choroid plexuses (CPs) and circumventricular organs (CVOs), demonstrating capillary-type-selective vascularization mechanisms. We found that zebrafish deficient for Gpr124, Reck, or Wnt7aa exhibit severely-impaired BBB angiogenesis without any apparent defect in fenestrated capillary formation in the CPs and CVOs. Conversely, simultaneous genetic loss of various Vegf combinations revealed remarkable heterogeneity of endothelial requirements for Vegfs-dependent angiogenesis within and across these organs, identifying unexpected interplay of Vegfc/d and Vegfa in fenestrated brain capillary formation. Expression analysis and paracrine activity-deficient *vegfc* mutant characterization suggest that endothelial cells and non-neuronal specialized cell types present in the CPs and CVOs are major sources of Vegfs responsible for regionally-restricted angiogenic interplay. Thus, local presentations and interplay of Vegfc/d and Vegfa control brain region-specific emergence of fenestrated capillaries, providing insight into fenestrated capillary formation in other organs and also how intra-organ vascular heterogeneity arises.

## INTRODUCTION

Throughout the body, the vascular system delivers oxygen, nutrients, and hormones while removing metabolic wastes to maintain homeostasis. In addition to this general function, each organ acquires unique vascular properties adapted to mediating its specific function during development, leading to the generation of vascular phenotypic heterogeneity in the organism (*1*). This inter-organ vascular heterogeneity is an evolutionarily conserved feature of the endothelium across diverse vertebrate species (*2*) and is important to facilitate timely physiological actions in response to constantly changing internal states in the organism. Recent bulk and single-cell RNA sequencing (scRNA-seq) has further validated significant transcriptional heterogeneity of endothelial cells across various organs in humans and mice (*3*–*6*) and even within the same tissue (*6*–*8*). How inter- and intra-organ endothelial heterogeneity is established and maintained is an outstanding question in vascular biology and organ physiology.

The brain exhibits striking intra-organ endothelial phenotypic heterogeneity (*6*, *8*–*10*). In most regions of the brain, endothelial cells form the semi-permeable blood-brain barrier (BBB), while those in the restricted regions of the brain develop highly permeable pores, or fenestrae. The BBB is an integral part of the neurovascular unit and forms a brain-vascular interface, which is vital for maintaining the optimal chemical and cellular milieu of the brain. In contrast, the choroid plexuses (CPs) and circumventricular organs (CVOs) are midline brain ventricular structures vascularized with fenestrated capillaries that lack the BBB. Fenestrated vasculature in these organs allow for high vascular permeability which serves to maintain fluid balance and also mediate neuroendocrine and neural immune activities (*7*, *11*). Thus, intra-brain vascular heterogeneity is fundamental to brain homeostasis and brain region-specific neural tasks. Additionally, under healthy conditions, CPs serve as a gateway for immune cells from the circulatory system to the brain. However, in neuroinflammatory conditions where BBB breakdown occurs, immune cells can penetrate the brain through vasculature that loses tight barrier properties, exacerbating neuroinflammatory responses. Thus, heterogeneous vascular barrier integrity is an important regulator of neural immune responses associated with age, injury, and/or disease. Despite the physiological and clinical importance, the mechanisms underlying intra-brain vascular heterogeneity establishment remain unclear. Moreover, the cellular and molecular basis of fenestrated brain capillary development remains poorly understood.

We recently reported that a unique set of angiogenic cues (Vegfab, Vegfc, and Vegfd) is required for fenestrated vessel formation in the zebrafish myelencephalic CP, an equivalent to the 4th ventricular CP in mammals (*12*). Intriguingly, the combined loss of these Vegfs has little impact on the formation of neighboring brain vasculature which displays BBB molecular signatures (*12*), indicating that endothelial cell-type-specific angiogenesis is a key mechanism driving local vascular heterogeneity within the brain. To further explore this possibility, we expanded our analysis to different CPs and CVOs in the present study to address the following questions. First, we asked whether Wnt/β-catenin signaling, a critical signaling pathway for BBB vascular development, plays a role in fenestrated brain vessel formation. Second, we investigated whether the combination of the Vegf ligands identified in our previous study is a universal angiogenic inducer of fenestrated vasculature across the central nervous system (CNS). Our combined analysis revealed fenestrated capillary type-selective angiogenic mechanisms, their local and inter-tissue regulatory heterogeneity, and a previously unappreciated function of Vegfc/d in this process.

## RESULTS

### Genetic loss of Wnt/β-catenin signaling leads to severely impaired angiogenesis in the brain parenchyma without any apparent defect in fenestrated brain vessel formation

Wnt/β-catenin signaling is well established as a central regulator of brain angiogenesis and BBB formation/maintenance (*13*, *14*). We first asked whether this signaling pathway is required for fenestrated vascular development across the CNS. We employed the zebrafish model due to its *ex-utero* development, rapid formation of brain vascular heterogeneity, and its facile visualization in whole-brain tissues of live animals. For example, 3D confocal imaging of the endothelial cell-specific transgenic (Tg) zebrafish reporter *Tg(kdrl:ras-mCherry)* carrying the *Et(cp:EGFP)* enhancer trap line allowed us to simultaneously visualize brain vasculature and two anatomically separate CPs, the diencephalic and myelencephalic CP (dCP and mCP) that are equivalent to the 3rd and 4th ventricular CP in mammals, respectively (Fig. 1A, 1A’) (*15*–*17*). Zebrafish larvae carrying double transgenic *Tg(plvap:EGFP);Tg(glut1b:mCherry)* reporters, which mark fenestrated endothelial cells in EGFP and BBB endothelium in mCherry, enabled us to map transcriptionally heterogeneous networks of brain and meningeal vasculature at 120 hours post fertilization (hpf) (Fig. 1B,1B’): Specifically, 1) blood vessels exhibiting high levels of *Tg(plvap:EGFP)*, but low levels of *Tg(glut1b:mCherry)*, expression; 2) those exhibiting low levels of *Tg(plvap:EGFP)*, but high levels of *Tg(glut1b:mCherry)*, expression; and 3) those with medium levels of both *Tg(plvap:EGFP)* and *Tg(glut1b:mCherry)* expression. As previously shown, vasculature that forms in tissues around the two CPs displays fenestrated molecular signatures represented by strong *Tg(plvap:EGFP)* expression (Fig. 1B, 1B’).

**Figure 1.**
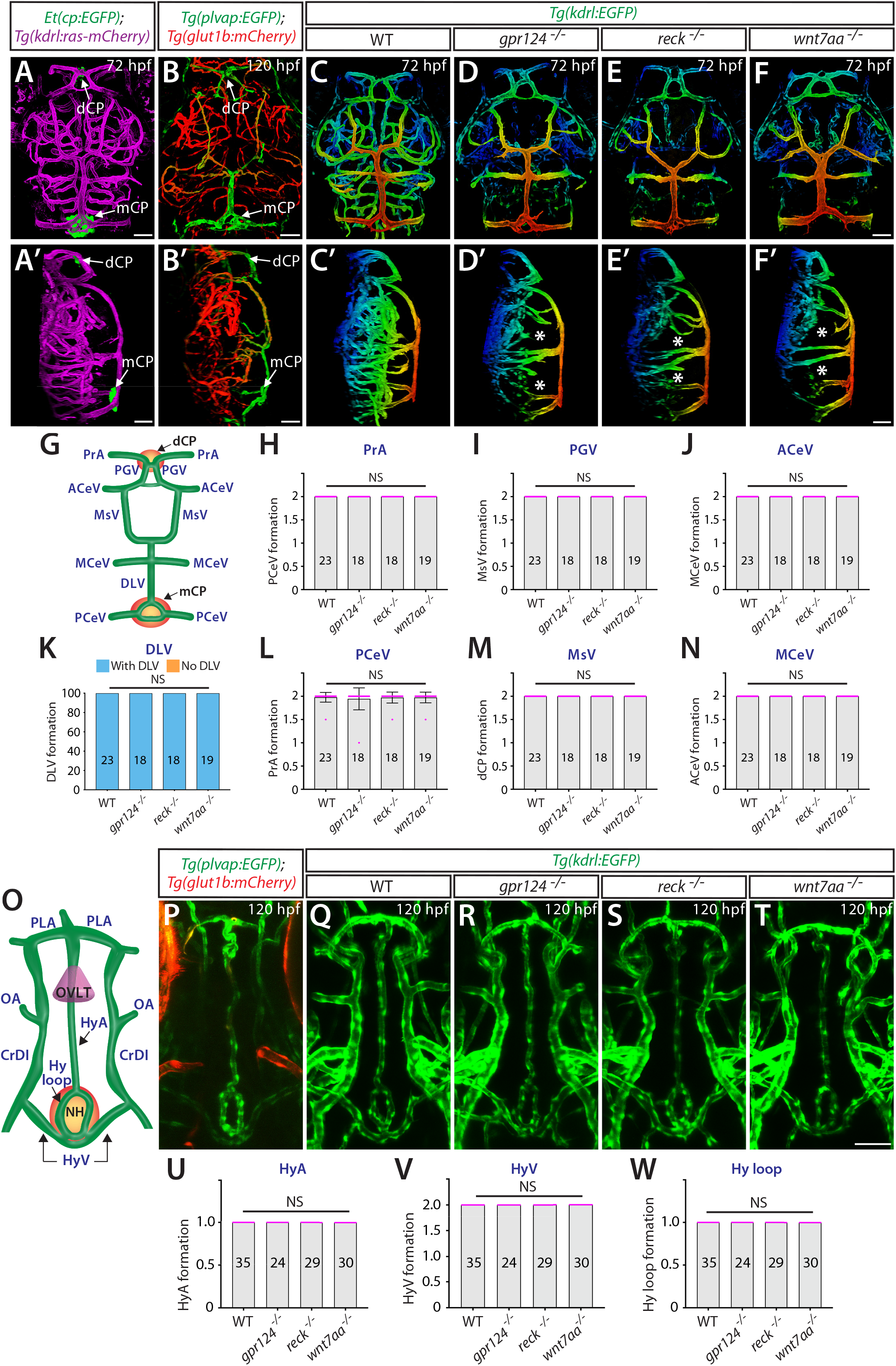
Genetic loss of Wnt/β-catenin signaling leads to severely impaired angiogenesis in the brain parenchyma without any apparent defect in fenestrated brain vessel formation. (**A**, **A’**) Dorsal (**A**) and lateral (**A’**) views of a 72 hpf *Et(cp:EGFP);Tg(kdrl:ras-mCherry)* larval head point to the locations of the diencephalic and myelencephalic CP (**dCP** and **mCP**, respectively). (**B**, **B’**) Dorsal (**B**) and lateral (**B’**) views of a 72 hpf *Tg(plvap:EGFP);Tg(glut1b:mCherry)* zebrafish head indicate fenestrated and BBB states of meningeal and brain vasculature. Blood vessels formed in the dCP and mCP display strong *Tg(plvap:EGFP)* fenestrated marker expression. (**C**–**F’**) Dorsal (**C**–**F**) and lateral (**C’**–**F’**) views of 72 hpf wild-type (WT) (**C**, **C’**), *gpr124*^-/-^ (**D**, **D’**), *reck*^-/-^ (**E**, **E’**), and *wnt7aa*^-/-^ (**F**, **F’**) zebrafish head vasculature visualized by *Tg(kdrl:EGFP)* expression. Color-coded maximum projection images indicate the most dorsal vessels in red and ventral ones in blue with a gradual color shift from dorsal to ventral. Asterisks indicate severely impaired angiogenesis in the brain parenchyma of these mutants compared to WT (**C’**–**F’**). (**G**) Schematic diagram of the dorsal view of cranial vasculature, illustrating the locations of the two CPs and distinct cranial blood vessels used for quantifications. **PrA**: prosencephalic artery, **PGV**: pineal gland vessel, **ACeV**: anterior cerebral vein, **MsV**: mesencephalic vein, **MCeV**: middle cerebral vein, **DLV**: dorsal longitudinal vein, **PCeV**: posterior cerebral vein. (**H**–**N**) Quantification of vessel formation in the dorsal meningeal and brain compartments at 72 hpf (n=23 for WT, n=18 for *gpr124*^-/-^, n=18 for *reck*^-/-^, and n=19 for *wnt7aa*^-/-^ fish). No significant difference was detected in *gpr124*^-/-^, *reck*^-/-^, or *wnt7aa*^-/-^ larvae compared to WT. (**O**) Schematic diagram of vasculature in the ventral diencephalon, illustrating the locations of the organum vasculosum of the lamina terminalis (**OVLT**), the neurohypophysis (**NH**), and distinct blood vessels used for quantifications. **HyA**: hypophyseal artery, **HyV**: hypophyseal veins, **Hy loop**: hypophyseal loop, **PLA**: palatocerebral arteries, **OA**: optic artery, **CrDI**: cranial division of the internal carotid artery. (**P**) Dorsal view of 120 hpf *Tg(plvap:EGFP);Tg(glut1b:mCherry)* ventral head shows strong *Tg(plvap:EGFP)* expression in the Hy loop, HyA, and PLA compared to its fainter signals in the HyV. (**Q**–**T**) Dorsal views of 120 hpf WT (**Q**), *gpr124*^-/-^ (**R**), *reck*^-/-^ (**S**), and *wnt7aa*^-/-^ (**T**) ventral head vasculature visualized by *Tg(kdrl:*EGFP) expression. Hypophyseal and OVLT vasculature forms in *gpr124*^-/-^, *reck*^-/-^, or *wnt7aa*^-/-^ larvae similar to WT. (**U**–**W**) Quantification of hypophyseal and OVLT vessel formation at 120 hpf (n=35 for WT, n=24 for *gpr124*^-/-^, n=29 for *reck*^-/-^, and n=30 for *wnt7aa*^-/-^ fish). No significant difference was detected in *gpr124*^-/-^, *reck*^-/-^, or *wnt7aa*^-/-^ larvae compared to WT. Data are means ± SD. NS: not significant. Scale bars: 50 μm in **A**–**B’**, in **F** for **C**–**F**, in **F’** for **C’**–**F’**, and in **T** for **P**–**T**.

To investigate the role of Wnt/β-catenin signaling in fenestrated brain vessel formation, we chose to analyze zebrafish that harbor mutations in *gpr124*, *reck*, or *wnt7aa*. Gpr124 and Reck act as co-activators of Wnt7a/b-specific canonical β-catenin pathway (*18*, *19*). Both *gpr124 ^s984^* and *wnt7aa ^ulb2^* zebrafish mutants carry out-of-frame mutations that induce a premature stop codon leading to truncated proteins that lack functional domains (*19*, *20*). The newly generated *reck^ulb3^* zebrafish mutants harbor a 12 bp in-frame deletion in the 3rd cysteine knot motif (Figure 1–figure supplement 1), close to the binding motifs of Reck to Gpr124 and Wnt7a/b that are critical for brain angiogenesis activity. We observed that *gpr124 ^s984^, reck^ulb3^*, or *wnt7aa ^ulb2^* mutants all displayed severe angiogenesis defects in the brain parenchyma (Fig. 1C–F”) as previously reported (*19*–*21*) and also similar to mouse knockouts (*22*–*24*), showing the high conservation of this signaling pathway in brain angiogenesis between zebrafish and mammals. Despite severe defects in brain parenchyma angiogenesis, we observed that meningeal angiogenesis and vascularization of both the dCP and mCP were not compromised in any of these mutants (Fig. 1G–N). In addition, we examined fenestrated vessel formation in the hypophysis and the organum vasculosum of the lamina terminalis (OVLT) (Fig. 1O), and observed no apparent vessel defect in these brain regions (Fig. 1P–W), indicating that fenestrated vessel development does not rely on Wnt/β-catenin signaling across the brain.

### Fenestrated vascular development at the interface of the dCP and pineal gland

We next sought to explore angiogenic cues responsible for vessel development in the fenestrated brain vascular beds, which is independent of Wnt/β-catenin signaling. Previous studies, including our recent work, characterized the vascularization processes of the mCP (*12*, *15*, *17*), however, no angiogenic cues have been reported for dCP vascularization. Intriguingly, through our literature reviews and immunostaining of *Tg(kdrl:ras-mCherry);Et(cp:EGFP)* larvae with an antibody for rhodopsin, a marker for pineal photoreceptor cells, we found that the dCP and pineal gland (PG) lie adjacent to vasculature (Fig. 2A–C”, Supplemental Movie 1) that displays strong fenestrated marker expression (Fig. 2D). The PG is the major neuroendocrine organ that secretes melatonin, which controls circadian rhythms and sleep-wake cycles (*25*). Identification of this unique dCP/PG vascular interface motivated us to determine its angiogenic mechanisms. The *Et(cp:EGFP)^+^* cells in the dCP were outlined by Claudin-5 tight junction protein expression, a marker for CP epithelial cells, allowing us to visualize dCP epithelial cells by immunostaining for Claudin-5 without the *Et(cp:EGFP)* reporter (Fig. 2E–E”).

**Figure 2.**
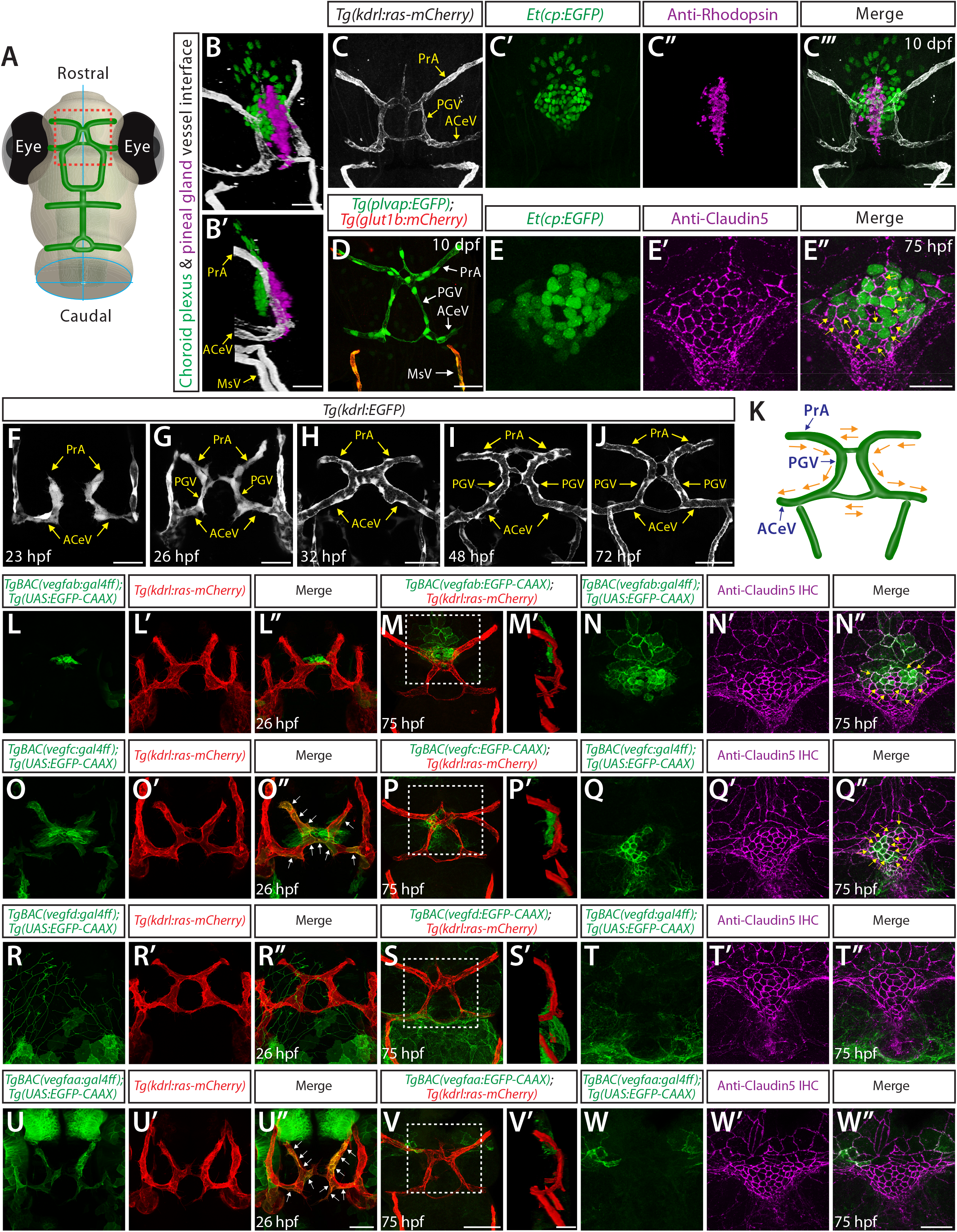
Fenestrated vascular development at the interface of the pineal gland and dCP, and *vegf* expression patterns during the development of this interface. (**A**) Schematic representation of the dorsal view of the zebrafish larval head, indicating the location of the dCP and pineal gland (**PG**) vascular interface with the red, boxed area. (**B**–**C”’**) 3D (**B**), lateral (**B’**), and dorsal (**C**–**C”’**) views of the 10 dpf *Et(cp:EGFP);Tg(kdrl:ras-mCherry)* head immunostained for rhodopsin show the 3D spatial relationship between dCP epithelial cells (green), pineal photoreceptor cells (magenta), and blood vessels (white). (**D**) Dorsal view of 10 dpf *Tg(plvap:EGFP);Tg(glut1b:mCherry)* zebrafish head shows strong *Tg(plvap:EGFP)* and undetectable *Tg(glut1b:mCherry)* expression shared in the PrA, PGV, and ACeV, which form in close proximity to the dCP/PG interface. In contrast, their neighboring vessels, MsV, display a different phenotype, including strong *Tg(glut1b:mCherry)* expression. (**E**–**E”**) Dorsal views of the 75 hpf *Et(cp:EGFP)* head immunostained for Claudin5 show EGFP^+^ dCP epithelial cells (yellow arrows) outlined by the tight junction protein Claudin5. (**F**–**J**) Dorsal views of 23 (**F**), 26 (**G**), 32 (**H**), 48 (**I**), and 72 (**J**) hpf *Tg(kdrl:EGFP)* rostral cranial vasculature show the developmental time courses of vascularization at the dCP/PG interface. (**K**) Schematic diagram of the vasculature shows the direction of blood flow at the dCP/PG interface. (**L**–**L”**) Dorsal views of a 26 hpf *TgBAC(vegfab:EGFP);Tg(kdrl:ras-mCherry)* embryonic head show *TgBAC(vegfab:EGFP)^+^* cells at the midline where bilateral PrA connect. (**M**–**N”**) Dorsal (**M**, **N**–**N”**) and lateral (**N’**) views of a 75 hpf *TgBAC(vegfab:EGFP);Tg(kdrl:ras-mCherry)* larval head immunostained for Claudin5. As compared to 26 hpf, an increased number of *TgBAC(vegfab:EGFP* ^+^ cells was observed at the PrA connection site and in its anterior brain regions (**M**, **M’**). Magnified images of the boxed area in (**M**) indicate *TgBAC(vegfab:EGFP)^+^* and Claudin5^+^ dCP epithelial cells (yellow arrows, **N**–**N”**). (**O**–**O”**) Dorsal views of a 26 hpf *TgBAC(vegfc:EGFP);Tg(kdrl:ras-mCherry)* embryonic head show *TgBAC(vegfc:EGFP)^+^* endothelial cells (white arrows) and separate cells at the PrA connection site. (**P**–**Q”**) Dorsal (**P**, **Q**–**Q”**) and lateral (**P’**) views of a 75 hpf *TgBAC(vegfc:EGFP);Tg(kdrl:ras-mCherry)* larval head immunostained for Claudin5. *TgBAC(vegfc:EGFP)^+^* cells were observed at the PrA connection site and in its posterior brain regions (**P, P’**). Magnified images of the boxed area in (**P**) indicate *TgBAC(vegfc:EGFP)^+^* and Claudin5^+^ dCP epithelial cells (yellow arrows, **Q**–**Q”**). (**R**–**R”**) Dorsal views of a 26 hpf *TgBAC(vegfd:EGFP);Tg(kdrl:ras-mCherry)* embryonic head show *TgBAC(vegfd:EGFP)^+^* meningeal fibroblast-like cells that reside posterior to the dCP/PG interface. *TgBAC(vegfd:EGFP)^+^* axonal projections were also visualized. (**S**–**T”**) Dorsal (**S, T**–**T”**) and lateral (**S’**) views of a 75 hpf *TgBAC(vegfd:EGFP);Tg(kdrl:ras-mCherry)* larval head immunostained for Claudin5. *TgBAC(vegfd:*EGFP)^+^ cells were observed in meningeal fibroblast-like cells that lie posterior to the PGV/ACeV (**S, S’**). Magnified images of the boxed area in (**s**) show no obvious *TgBAC(vegfd:*EGFP) expression in dCP epithelial cells (**T**–**T”**). (**U**–**U”**) Dorsal views of a 26 hpf *TgBAC(vegfaa:EGFP);Tg(kdrl:ras-mCherry)* embryonic head show *TgBAC(vegfaa:EGFP)^+^* endothelial cells (white arrows). (**V**–**W”**) Dorsal (**V**, **W**–**W”**) and lateral (**V’**) views of a 75 hpf *TgBAC(vegfaa:EGFP);Tg(kdrl:ras-mCherry)* larval head immunostained for Claudin5. Sparse *TgBAC(vegfaa:EGFP)^+^* cells were observed at the lateral periphery of the dCP (**W**, **W’**). Magnified images of the boxed area in (**V**) show *TgBAC(vegfaa:*EGFP)^+^ and Claudin5^+^ dCP epithelial cells at the periphery. Scale bars: 30 μm in **B**, **B’**, **D**, in **C”’** for **C**–**C”’**, in **E”** for **E**–**E”**, in **V’** for **M’**, **P’**, **S’**; 50 μm in **F**–**J**, in **U”** for **L**–**L”**, **O**–**O”**, **R**–**R”**, **U**–**U”**, in **V** for **M**, **P**, **S**, in **W”** for **N**–**N”**, **Q**–**Q”**, **T**–**T”**, **W**–**W”**.

Angiogenic steps leading to vascularization of the dCP/PG interface have not been well characterized. We noted that this vascularization process initiates at very early embryonic stages. By 23 hpf, the anterior cerebral vein (ACeV) sprouts bilaterally from the primordial hindbrain channel and extends dorsally to start forming the bilateral prosencephalic artery (PrA) (Fig. 2F). The PrA then extends anteroventrally to connect with the cranial division of the internal carotid artery around 26 hpf (Fig. 2G). Anastomosis of this vascular plexus occurs by 32 hpf (Fig. 2H) after the systemic circulation of blood begins at approximately 24-26 hpf (*26*). Through 72 hpf, the vascular structure undergoes remodeling and maturation to establish a functional circuit (Fig. 2I, 2J). Close examination of blood circulation under phase-contrast imaging reveals directional blood circulation from the PrA through the ACeV via the PG vessel (PGV) which we term to specify (Fig. 2K).

### BAC transgenic analysis of *vegf* expression at the developing dCP/PG interface

Since no angiogenic mechanisms have been reported with respect to the vascularization of the dCP/PG interface, we first explored where and which *vegfs* are expressed during the development of this interface. To visualize the expression of individual *vegf* isoforms at the single-cell resolution, we employed our recently generated BAC transgenic Gal4FF reporter lines that reliably recapitulate each of the endogenous gene expression detected by *in situ* hybridization (*12*, *27*). The individual Gal4FF drivers were first crossed with *Tg(UAS:EGFP-CAAX)* fish, which carry a membrane-bound EGFP gene downstream of upstream activation sequence (UAS). The resultant double Tg fish are here abbreviated *TgBAC(vegfab:EGFP)*, *TgBAC(vegfc:EGFP)*, *TgBAC(vegfd:EGFP)*, and *TgBAC(vegfaa:EGFP)*. All of these Tg lines were further crossed with the endothelial *Tg(kdrl:ras-mCherry)* reporter to visualize *vegf*-expressing cells and vascular endothelial cells simultaneously. Using these Tg lines, we examined spatial relationships between *vegf*-expressing cells and forming vasculature.

We noted that at 26 hpf, *TgBAC(vegfab:EGFP)* expression is specifically localized at the junction where bilateral PrA joins (Fig. 2L–L”). At 75 hpf, an increased number of *TgBAC(vegfab:EGFP)^+^* cells were seen in this region (Fig. 2M), which were co-labelled with an antibody for Claudin-5, a marker for dCP epithelial cells (Fig. 2E–E”, 2N–N”). On the other hand, we observed the broader and prominent *TgBAC(vegfc:EGFP)* expression in two cell types around this region at 26 hpf (Fig. 2O–O”): 1) developing endothelial cells that are part of the ACeV, PGV, and PrA and 2) cells at the junction where bilateral PrA joins. By 75 hpf, *TgBAC(vegfc:EGFP)* expression in endothelial cells disappeared (Fig. 2P), but *TgBAC(vegfc:EGFP)^+^* cells marked by Claudin-5 immunoreactivity were still observed (Fig. 2Q–Q”). These *TgBAC(vegfc*:EGFP)^+^ cells lie at the PrA junction slightly posterior to where *TgBAC(vegfab*:EGFP) expression was seen (Fig. 2M’, 2P’). In contrast, *TgBAC(vegfd:EGFP)* expression was not observed at the PrA junction or in dCP epithelial cells at 26 and 75 hpf (Fig. 2R–T”). *TgBAC(vegfaa:EGFP)* expression was found in developing endothelial cells that are part of the ACeV, PGV, and PrA at 26 hpf (Fig. 2U–U”). This endothelial *TgBAC(vegfaa:EGFP)* expression disappeared by 75 hpf (Fig. 2V, 2V’), and only sparsely labelled cells were observed in the region close to the dCP (Fig. 2W–W”).

To summarize, *TgBAC(vegfc:EGFP)* and *TgBAC(vegfaa:EGFP)* expression was observed similarly in extending endothelial cells at 26 hpf, which disappeared by 75 hpf. *TgBAC(vegfc:EGFP)* and *TgBAC(vegfab:EGFP)* expression at 26 hpf appears to mark cells that subsequently differentiate and/or maturate into dCP epithelial cells by 75 hpf. These overlapping expression patterns imply potential functional redundancy among multiple Vegf ligands in regulating vascularization at the dCP/PG interface.

### Anatomically separate CPs require a distinct set of Vegf ligands for vascularization

We previously illustrated that multiple Vegf ligands (Vegfab, Vegfc, and Vegfd) are redundantly involved in the development of fenestrated mCP vasculature (*12*). To determine whether this molecular combination universally regulates fenestrated vessel formation across the brain, we first asked whether the same set of Vegf ligands is required for the vascularization of dCP/pineal gland interface. To this aim, we analyzed *vegfab^bns92^, vegfc^hu6410^*, and *vegfd^bns257^* mutants that carried the endothelial *Tg(kdrl:EGFP)* reporter individually and in all possible combinations. We chose to analyze these mutants at 10 days post fertilization (dpf) – a stage when most of the major blood vessels are formed (*26*) and that is a week after vasculature at the dCP/PG interface normally forms. Conducting a phenotypic analysis at this stage allowed us to eliminate the possibility of developmental delays in mutants.

At 10 dpf, none of the *vegfab^bns92^, vegfc^hu6410^*, and *vegfd^bns257^* single mutants displayed a drastic difference compared to WT, except a mild, yet significant, defect in PrA formation in *vegfc^hu6410^* mutants (Fig. 3A–D, 3I). The combined loss of the three genes led to significantly exacerbated phenotypes in fenestrated vessel formation at the dCP/PG interface (Fig. 3H). The most pronounced defect observed was on PrA formation, although this phenotype is partially penetrant (Fig. 3I). One-third of the triple mutants lacked bilaterally-formed PrA, 40% triple mutants exhibited only unilateral PrA, and only 20% mutants formed bilateral PrA. ACeV formation deficit is milder in triple mutants, with approximately 27% of them exhibiting unilateral ACeV. Statistical analysis of these genetic data shows that *vegfab* and *vegfc* genetically interact in PrA formation with little effect on dCP and ACeV formation. In contrast, *vegfd* and *vegfc* genetically interact in regulating dCP and ACeV, but not PrA, formation. While these results indicate the requirements of all the three Vegf ligands for vascularization of this interface, the phenotypes observed in triple mutants are much milder than those observed in the mCP where we found a near-complete penetrance and loss of fenestrated vasculature (*12*). These data imply differential molecular requirements for fenestrated angiogenesis across the CPs.

**Figure 3.**
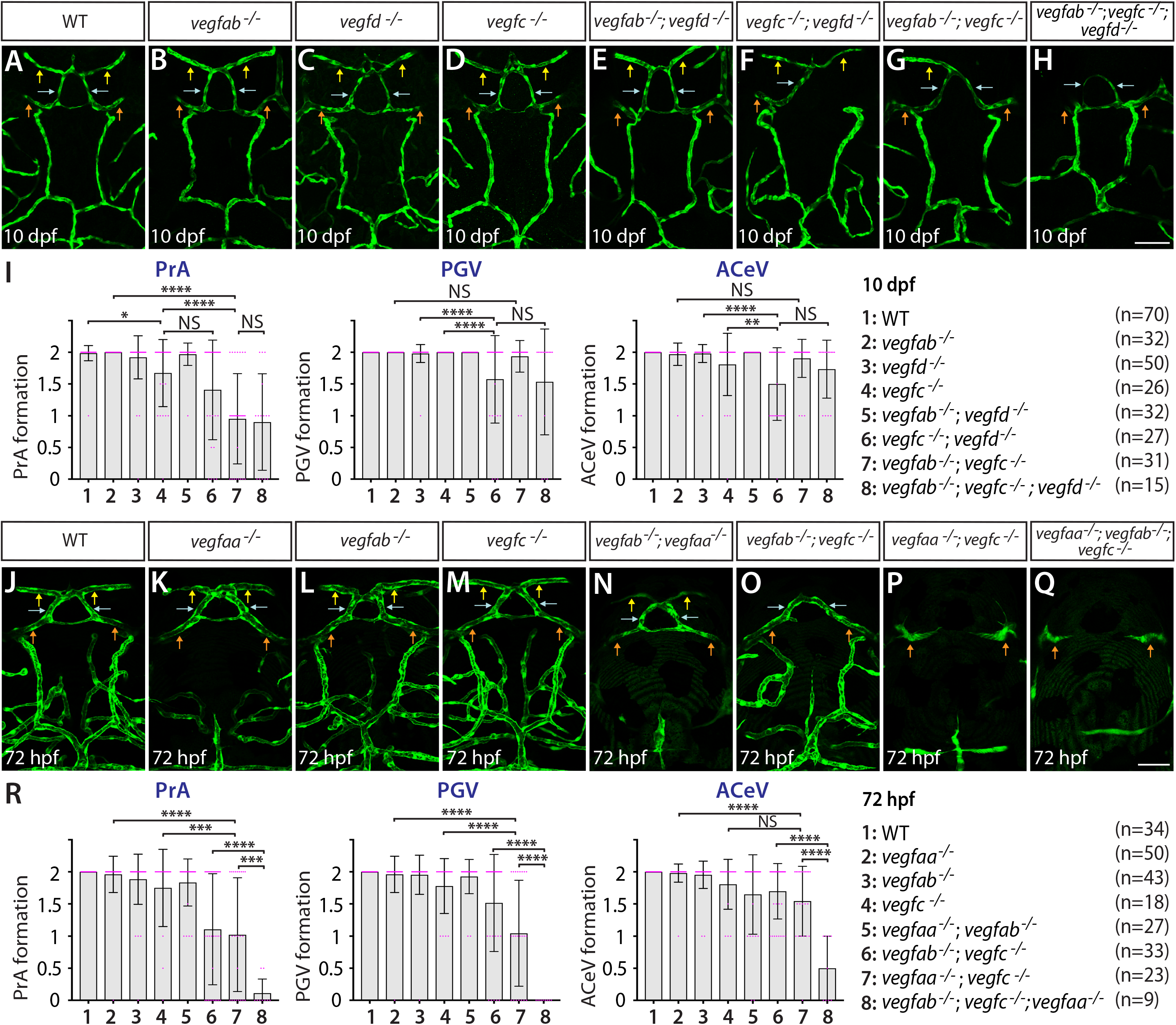
Local regulatory heterogeneity of Vegfs-dependent angiogenesis at the dCP/PG interface. (**A**–**H**) Dorsal views of 10 dpf WT (**A**), *vegfab*^-/-^ (**B**), *vegfd*^-/-^ (**C**), *vegfc*^-/-^ (**D**), *vegfab^-/-^;vegfd^-/-^* (**E**), *vegfc^-/-^;vegfd^-/-^* (**F**), *vegfab^-/-^;vegfc^-/-^* (**G**), and *vegfab^-/-^;vegfc^-/-^;vegfd^-/-^* (**H**) cranial vasculature visualized by *Tg(kdrl*:EGFP) expression. Yellow arrows point to the PrA, blue arrows to the PGV, and orange arrows to the ACeV. A majority of *vegfab^-/-^;vegfc^-/-^* (**G**) and *vegfab^-/-^;vegfc^-/-^;vegfd^-/-^* (**H**) larvae lacked the PrA at either or both sides. *vegfc^-/-^;vegfd^-/-^* (**F**) and *vegfab^-/-^;vegfc^-/-^;vegfd^-/-^* (**H**), but not *vegfab^-/-^;vegfc^-/-^* (**G**), larvae displayed partially penetrant defects in PGV and/or ACeV formation. (**I**) Quantification of PrA, PGV, and ACeV formation at 10 dpf (the number of animals examined per genotype is listed in the panel). Statistical data support genetic interactions between *vegfab* and *vegfc* in PrA formation and between *vegfd* and *vegfc* in PGV and ACeV formation. No significant contributions of *vegfd* or *vegfab* to the formation of the PrA or PGV/ACeV, respectively, were noted. (**J**–**Q**) Dorsal views of 72 hpf WT (**J**), *vegfaa*^-/-^ (**K**), *vegfab*^-/-^ (**L**), *vegfc*^-/-^ (**M**), *vegfab^-/-^;vegfaa^-/-^* (**N**), *vegfab^-/-^;vegfc^-/-^* (**O**), *vegfaa^-/-^;vegfc^-/-^* (**P**), and *vegfaa^-/-^;vegfab^-/-^;vegfc^-/-^* (**Q**) cranial vasculature visualized by *Tg(kdrl*:EGFP) expression. Yellow arrows point to the PrA, blue arrows to the PGV, and orange arrows to the ACeV. *vegfab^-/-^;vegfc^-/-^*, but not their respective single mutants, exhibited pronounced PrA formation deficits. *vegfaa^-/-^* and *vegfab^-/-^;vegfaa^-/-^* displayed severe defects in MsV formation without a deficit in PrA, PGV, or ACeV development. *vegfaa^-/-^;vegfc^-/-^* and *vegfaa^-/-^;vegfab^-/-^;vegfc^-/-^* larvae exhibited a severe loss of the PrA and PGV. (**R**) Quantification of PrA, PGV, and ACeV formation at 72 hpf (the number of animals examined per genotype is listed in the panel). Statistical data support genetic interactions between *vegfab* and *vegfc* in PrA formation and between *vegfaa* and *vegfc* in PrA and PGV formation. Furthermore, significant genetic interactions were detected among these three genes in vascular formation at this interface. Scale bars: 50 μm in **H** for **A**–**H** and in **Q** for **J**–**Q**.

To determine additional angiogenic factor(s) critical for vascularization of the dCP/PG interface, we analyzed zebrafish mutants that lack another Vegfa paralog, Vegfaa. Since *vegfaa^bns1^* mutants were previously shown to die at approximately 5 dpf due to the severe early embryonic vascular defects (*28*), we analyzed this mutant at 72 hpf. *vegfaa^bns1^* single mutants did not show a defect in fenestrated vessel development at the dCP/PG interface (Fig. 3K, 3R). However, they did display a severe defect in the formation of the neighboring bilateral MsV that displays medium levels of both *Tg(plvap:EGFP)* and *Tg(glut1b:mCherry)* expression (Fig. 2D) and BBB marker Claudin5 immunoreactivity (*12*). When we deleted *vegfaa* and *vegfc* simultaneously, PrA and PGV formation defects were substantially enhanced (Fig. 3P). Statistical analysis supports a strong genetic interaction between *vegfc* and *vegfaa* in the development of the PrA and PGV, but not of the ACeV. The combined loss of *vegfab* and *vegfc* induced striking defects specifically in PrA formation similar to what was observed at 10 dpf (Fig. 3O). Intriguingly, the combined loss of *vegfaa* and *vegfab*, the two Vegfa paralogs in zebrafish, did not cause a defect in vessel formation at this interface (Fig. 3N, 3R).

Collectively, these mutant analyses uncover surprising regulatory heterogeneity of angiogenesis within the local brain environment. Specifically, the results reveal Vegfc’s genetic interactions with *vegfab, vegfd*, or *vegfaa* in regulating fenestrated vessel development at the dCP/PG interface, placing Vegfc as a central angiogenic regulator of this process. This angiogenic requirement is different from that needed for mCP vascularization which involves a combination of Vegfab, Vegfc, and Vegfd in the way that Vegfab acts as a central angiogenic factor. This distinction may be due to brain regional differences in CP molecular signatures, including secretome, as indicated by the recent transcriptomic studies of CPs dissected from different ventricles (*29*, *30*).

### Endothelial cell-autonomous and cell non-autonomous requirements of Vegfc for vascularization of the dCP/PG interface

Our expression and genetic data indicate that *vegfc* is expressed in both developing endothelial cells and dCP epithelial cells during dCP/PG vascularization and that Vegfc functionally interacts with other Vegfs in controlling this process. Indeed, we observed *vegfab* expression in developing dCP epithelial cells and *vegfaa* expression in extending endothelial cells during dCP/PG vascularization. These observations led us to hypothesize that Vegfc’s angiogenic activity involves both endothelial cell-autonomous and cell non-autonomous actions in this context mediated by the discrete Vegfa paralogs.

To test this hypothesis, we employed a separate *vegfc* mutation, *vegfc^um18^*, which was previously isolated from a forward genetic screen (*31*). This particular mutation was shown to generate a prematurely truncated Vegfc protein that lacks efficient secretory and paracrine activity but retains the ability to activate its Flt4 receptor (in another words, cell-autonomous activity of Vegfc is retained) (*31*). Homozygous embryos carrying this particular mutation alone did not exhibit a defect in dCP/PG vascularization at 72 hpf (Fig. 4C). However, we noted that the *vegfc^um18^* mutant allele displayed a genetic interaction with *vegfab*, leading to a partially penetrant, yet significant, defect in PrA formation (Fig. 4A–F). The extent of this PrA defect is milder than that was noted in double mutants of *vegfab;vegfc^hu6410^* larvae analyzed at the same stage (Fig. 4I). These results suggest that reduced Vegfc paracrine activity alone exhibits a milder defect than that observed in fish that lack both paracrine and autocrine Vegfc activities, indicating that endothelial cell-autonomous and cell non-autonomous Vegf activities are necessary for PrA formation.

**Figure 4.**
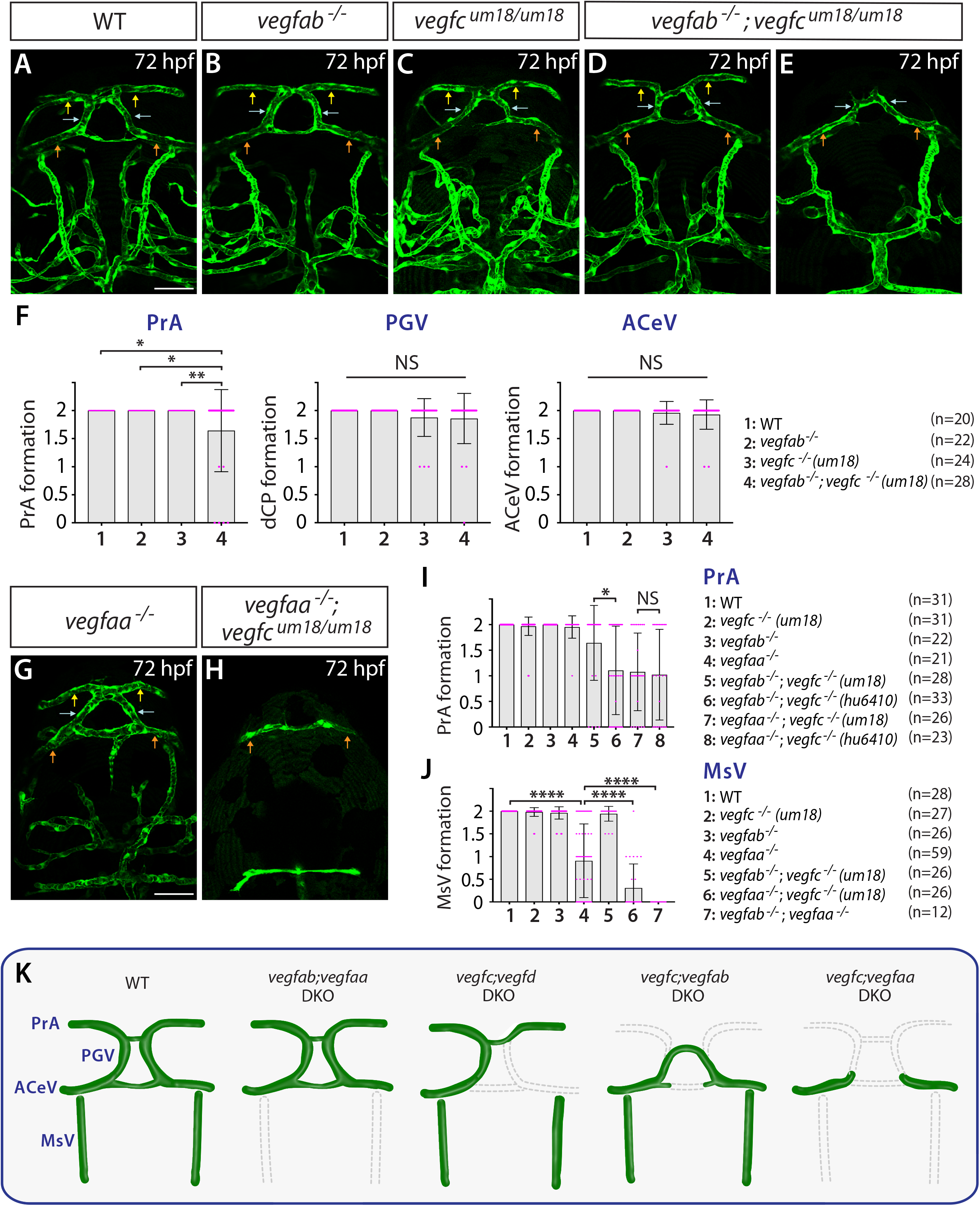
Endothelial cell-autonomous and cell non-autonomous requirements of Vegfc for vascularization of the dCP/PG interface. (**A**–**E**) Dorsal views of 72 hpf WT (**A**), *vegfab*^-/-^ (**B**), *vegfc^um18/um18^* (**C**), and *vegfab^-/-^;vegfc^um18/um18^* (**D**, **E**) cranial vasculature visualized by *Tg(kdrl*:EGFP) expression. Yellow arrows point to the PrA, blue arrows to the PGV, and orange arrows to the ACeV. Although none of *vegfab^-/-^* and *vegfc^um18/um18^* fish exhibited a defect in PrA formation (**B**, **C**), approximately 21% of *vegfab^-/-^;vegfc^um18/um18^* larvae lacked the PrA at either or both sides (**E**). (**F**) Quantification of PrA, PGV, and ACeV formation at 72 hpf (the number of animals examined per genotype is listed in the panel). Specific and significant defect was observed in PrA formation in *vegfab^-/-^;vegfc^um18/um18^* larvae compared to other three genotypes. (**G**, **H**) Dorsal views of 72 hpf *vegfaa*^-/-^ (**G**) and *vegfaa^-/-^;vegfc^um18/um18^* (**H**) cranial vasculature visualized by *Tg(kdrl*:EGFP) expression. Yellow arrows point to the PrA, blue arrows to the PGV, and orange arrows to the ACeV. Although *vegfaa^-/-^* or *vegfc^um18/um18^* larvae fully formed fenestrated vasculature at the dCP/PG interface, most of *vegfaa^-/-^;vegfc^um18/um18^* larvae failed to form the PrA and PGV at either or both sides. (**I**) Quantification of PrA formation at 72 hpf (the number of animals examined per genotype is listed in the panel). The quantitative results of several genotypes were presented again or integrated in this graph for comparison purposes. Previously presented results are the *vegfc^-/-^* (*hu6410* allele) related data from Fig. 3R, and the data in Fig. 4F were either re-presented or combined with the quantitative results shown in Fig. 4G, 4H. Paracrine activity-deficient *vegfc^um18/um18^* larvae in the *vegfab*^-/-^ background displayed a significantly milder defect in PrA formation than that observed in *vegfab^-/-^;vegfc^-/-^* (*hu6410* allele) animals that lack both endothelial cell-autonomous and cell non-autonomous Vegfc function. (**J**) Quantification of MsV formation at 72 hpf (the number of animals examined per genotype is listed in the panel). Severe defects in MsV formation in *vegfaa^-/-^* larvae were exacerbated by genetic deletions of *vegfc* (*um18* allele) or *vegfab*. (**K**) Schematic representations of the robust vascular phenotypes at the dCP/PG interface that were observed in various *vegf* mutant backgrounds. Genetic evidence suggests that there are specific Vegf requirements for the formation of individual blood vessels necessary for full vascularization of the dCP/PG interface. These results indicate highly heterogeneous requirements for angiogenesis within the local brain environment. Scale bars: 50 μm in **A** for **A**–**E** and in **G** for **G**–**H**.

We next analyzed double mutants of *vegfaa;vegfc ^um18^* larvae to gain additional insight into Vegfc angiogenic action. We found that *vegfaa;vegfc ^um18^* larvae exhibited a similar extent of the defect observed in *vegfaa;vegfc^hu6410^* larvae examined at the same stage (Fig. 4G–I), suggesting that a loss of Vegfc paracrine activity has a profound impact on PrA formation in *vegfaa* mutant background. We noted that the formation of neighboring blood vessels, MsV, which displays medium levels of both BBB and fenestrated marker expression, was severely abrogated in the absence of Vegfaa alone (Fig. 4G, 3K) and further exacerbated in combination with Vegfab or Vegfc deletion (Fig. 4H, 4J). These results show that Vegfaa plays a major role in regulating MsV development, providing additional evidence for regulatory heterogeneity of angiogenesis within this brain region. This local regulatory heterogeneity of angiogenesis is summarized in Fig. 4K.

### Normal choriocapillaris formation in zebrafish deficient for Wnt/β-catenin signaling, and conserved expression of Vegfa paralogs in retinal pigment epithelium

The choriocapillaris, or the choroidal vascular plexus (CVP), is a dense network of fenestrated capillaries that form in adjacent tissues outside of the retinal pigment epithelial cell (RPE) layer in the eyes. This dense vascular network mediates efficient exchanges of nutrients and metabolic wastes in the outer retina, including RPE and outer nuclear layers where photoreceptors are present. Previous studies in mice indicate that *Vegfa* is expressed in RPE and is critical for the formation of fenestrated choriocapillaris (*32*–*34*). In zebrafish, Vegfr2 receptor paralogs, Kdrl and Kdr, have been implicated to regulate choriocapillaris development (*35*), however, it remains unclear which Vegf ligands are involved in this process. Moreover, given the fact that this capillary development is better characterized than that in the CPs and CVOs in mammals, we expected that this model could help address conservation of *vegf* expression and function in fenestrated CNS angiogenesis between mammals and zebrafish.

Although *Tg(plvap:EGFP)* fenestrated marker expression is somewhat weak in the choriocapillaris at early larval stages, we noted endothelial fenestrations in this capillary network at the ultrastructure level (Fig. 5A–-C). Confocal visualization of the choriocapillaris from the back of isolated eyes reveals no apparent defect in choriocapillaris formation in *gpr124, reck*, or *wnt7aa* mutants at 6 dpf (Fig. 5D–I). These results suggest that Wnt/β-catenin signaling is not involved in fenestrated choriocapillaris development similarly to what is shown in fenestrated brain vascular beds (Fig. 1).

**Figure 5.**
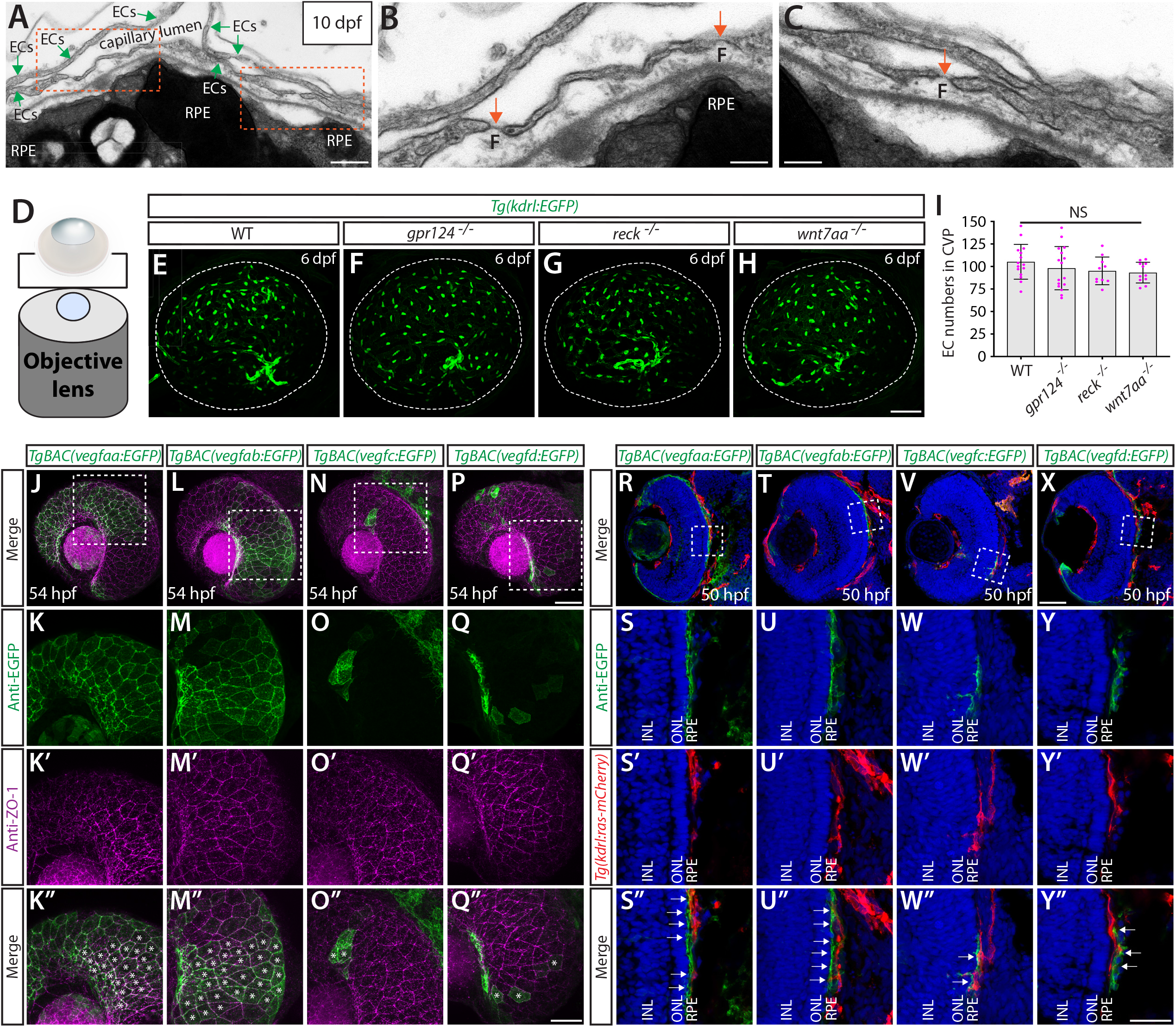
Normal choriocapillaris formation in zebrafish deficient for Wnt/β-catenin signaling, and conserved expression of Vegfa paralogs in retinal pigment epithelium. (**A**–**C**) Transmission electron microscopy images of 10 dpf WT outer retina focused on the choriocapillaris and retinal pigment epithelium (**RPE**) layer. Magnified images of the boxed areas in (**A**) show the presence of fenestrae (**F**) in the endothelial cells comprising the choriocapillaris (orange arrows, **B**, **C**). (**D**) Schematic diagram of 3D confocal choriocapillaris imaging from the back of dissected eyes. (**E**–**H**) 6 dpf WT (**E**), *gpr124^-/-^* (**F**), *reck^-/-^* (**G**), and *wnt7aa^-/-^* (**H**) choriocapillaris visualized by *Tg(kdrl:EGFP)* expression. Confocal *z*-stack images of dissected eyeballs were taken after immunostaining for GFP from the back of the eyeballs. (**I**) Quantification of endothelial cells that comprise the choriocapillaris at 6 dpf (n=15 for WT, n=16 for *gpr124^-/-^*, n=10 for *reck^-/-^*, and n=11 for *wnt7aa^-/-^* fish). No significant difference was observed across these genotypes. (**J**–**O”**) 54 hpf *TgBAC(vegfaa:EGFP)* (**J**), *TgBAC(vegfab:EGFP)* (**L**), *TgBAC(vegfc:EGFP)* (**N**), and *TgBAC(vegfd:EGFP)* (**P**) embryos immunostained for GFP and ZO-1, a tight junction marker for RPE. Magnified images of the boxed areas in (**J**), (**L**), (**N**), and (**P**) are shown in (**K**–**K”**), (**M**–**M”**), (**O**–**O”**), and (**Q**–**Q”**), respectively. *TgBAC(vegfaa:EGFP)* and *TgBAC(vegfab:EGFP)* expression was broadly co-localized with ZO-1 immunoreactive signals in RPE (asterisks in **K”, M”**). Sparse EGFP^+^ cells were observed in *TgBAC(vegfc:EGFP)* and *TgBAC(vegfd:EGFP)* embryonic eyes, some of which were co-localized with ZO-1 immunoreactivity (asterisks in **O”**, **Q”**). (**R**–**Y”**) Cryosections of 50 hpf *TgBAC(vegfaa:EGFP)* (**R**), *TgBAC(vegfab:EGFP)* (**T**), *TgBAC(vegfc:EGFP)* (**V**), and *TgBAC(vegfd:EGFP)* (**X**) embryos that carried the *Tg(kdrl:ras-mCherry)* endothelial transgene. Sections were immunostained for GFP and DsRed. Magnified images of the boxed areas in (**R**), (**T**), (**V**), and (**X**) are shown in (**S**–**S”**), (**U**–**U”**), (**W**–**W”**), and (**Y**–**Y”**), respectively. *TgBAC(vegfaa:EGFP)* and *TgBAC(vegfab:EGFP)* expression was broadly observed in the RPE layer directly adjacent to the choriocapillaris (white arrows in **S”**, **U”**). Sparse EGFP^+^ cells in *TgBAC(vegfc:EGFP)* and *TgBAC(vegfd:EGFP)* embryonic sections lied in close proximity to the choriocapillaris (white arrows in **W”**, **Y”**). **INL**: inner nuclear layer, **ONL**: outer nuclear layer. Scale bars: 500 nm in **A**; 200 nm in **B**, **C**; 50 μm in **H** for **E**–**H**, in **P** for **J**, **L**, **N**, and in **X** for **R**, **T**, **V**; 30 μm in **Q”** for **K**–**K”**, **M**–**M”**, **O**–**O”**, **Q**–**Q”**; 25 μm in **Y”** for **S**–**S”**, **U**–**U”**, **W**–**W”**, **Y**–**Y”**.

Next, we examined *vegf* expression using our BAC transgenic lines and found that at 54 hpf when active angiogenesis is occurring to form the choriocapillaris (*35*, *36*), both *vegfaa* and *vegfab* BAC reporter expression is co-labelled with an antibody for ZO-1, a tight junction protein marker for RPE (Fig. 5J–M”). Cryosections of these transgenic embryos at 50 hpf further confirmed strong and specific *vegfaa* and *vegfab* BAC reporter expression in the RPE layer within the outer retina, which lies adjacent to *Tg(kdrl:*ras-mCherry)^+^ endothelial cells comprising the choriocapillaris (Fig. 5R–U”). Restriction of *vegfaa* and *vegfab* expression to RPE is consistent with *Vegfa* expression patterns reported in developing murine retina (*32*, *33*). As compared to the broad expression of *vegfaa* and *vegfab* in RPE, *vegfc* and *vegfd* BAC reporter expression marked retinal cells only sparsely, and a vast majority of RPE cells were devoid of reporter expression (Fig. 5N–Q”, 5V–-Y”).

### Zebrafish Vegfa paralog functional redundancy in choriocapillaris formation supports high conservation of fenestrated CNS angiogenic mechanisms between zebrafish and mammals

To determine which Vegf ligands are involved in choriocapillaris development, we crossed mutants with *Tg(kdrl:EGFP):Tg(kdrl:NLS-mCherry)* double transgenic reporters that label endothelial cell bodies in EGFP and their nuclei in mCherry. These reporter lines enabled us to quantify the exact number of endothelial cells that comprise the choriocapillaris. Since a previous time-lapse study indicates that a vascular network of the choriocapillaris is mostly formed by around 65 hpf (*36*), we chose to analyze mutants at 72 hpf or later. Based on prominent *vegfaa* and *vegfab* BAC reporter expression in RPE, we first examined mutants of these two genes by incrossing double heterozygous adults and analyzing their progeny at 72 hpf. Individual homozygous mutants displayed a considerable reduction in endothelial cell numbers of the choriocapillaris, resulting in diminished vascular network elaboration, as compared to WT (Fig. 6A–C). Homozygous *vegfaa* mutants have underdeveloped eyes likely due to poor vascularization of the eye. Importantly, we observed a strong genetic interaction between these two Vegfa paralogs, with increased deletion of either gene resulting in reduced numbers of endothelial cells that comprise the choriocapillaris (Fig. 6A–E). This finding indicates that both Vegfa paralogs are required for fenestrated choriocapillaris development.

**Figure 6.**
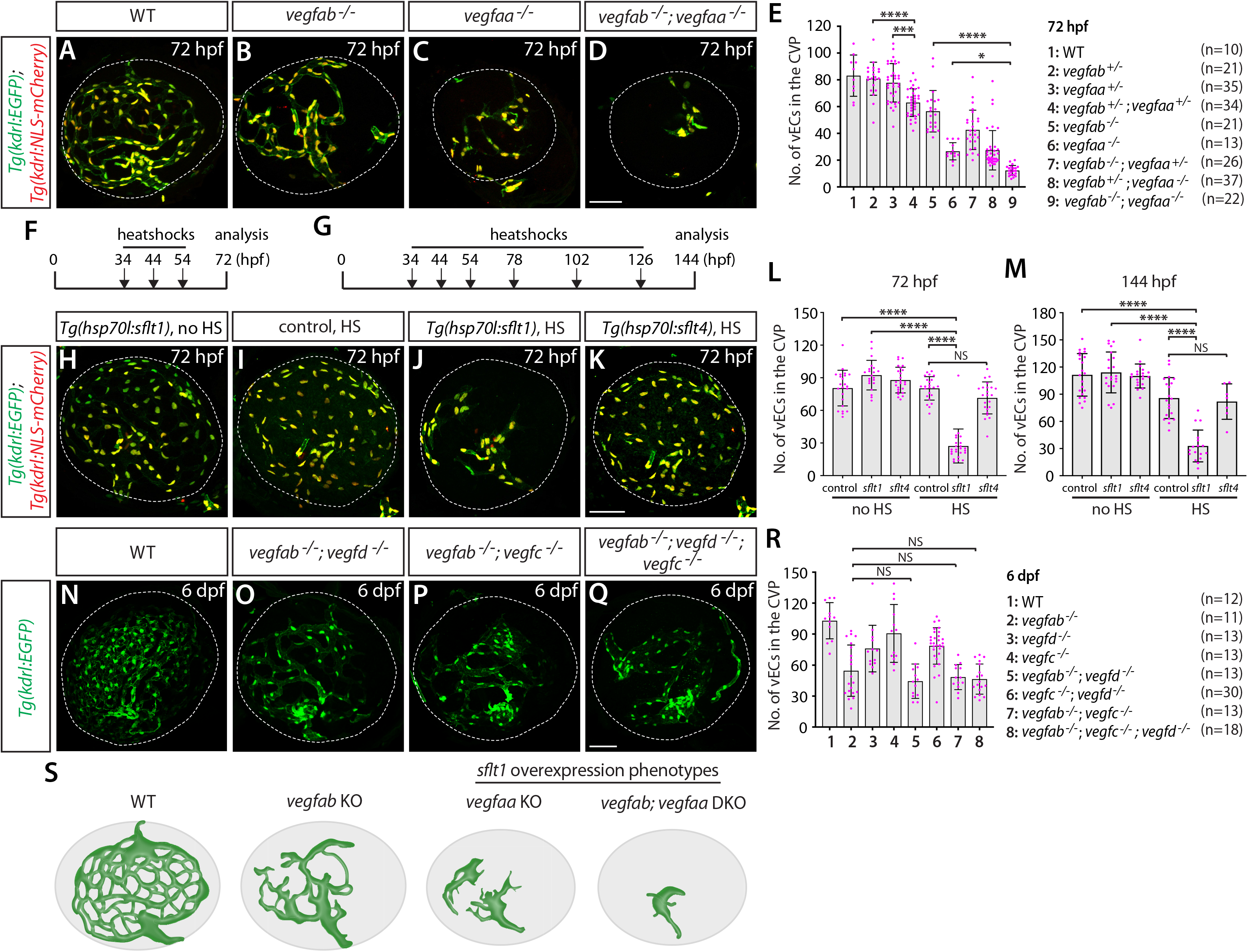
Zebrafish Vegfa paralogs redundantly regulate fenestrated choriocapillaris formation. (**A**–**D**) 72 hpf WT (**A**), *vegfab^-/-^* (**B**), *vegfaa^-/-^* (**C**), and *vegfab^-/-^;vegfaa^-/-^* (**D**) choriocapillaris (choroidal vascular plexus, **CVP**) visualized by *Tg(kdrl:EGFP)* and *Tg(kdrl:*NLS-mCherry) expression after immunostaining for GFP and DsRed. (**E**) Quantification of the number of endothelial cells that comprise the CVP at 72 hpf (the number of animals examined per genotype is listed in the panel). (**F**, **G**) Experimental time course of the heat-shock experiments for panels (**H**–**M**). *Tg(hsp70l:sflt1), Tg(hsp70l:sflt4)*, and their control siblings were subjected to a heat-shock at 34, 44, and 54 hpf for analysis at 72 hpf (**F**) or subjected to additional heatshocks at 78, 102, and 126 hpf for analysis at 144 hpf (**G**). (**H**–**K**) The CVP of 72 hpf *Tg(hsp70l:sflt1);Tg(kdrl:EGFP);Tg(kdrl:NLS-mCherry)* (**H**, **J**), *Tg(kdrl:EGFP);Tg(kdrl:NLS-mCherry)* (**I**) and *Tg(hsp70l:sflt4);Tg(kdrl:EGFP);Tg(kdrl:NLS-mCherry)* (**K**) larvae after treatment with (**I**–**K**) and without (**H**) multiple heatshocks. Heatshock-induced overexpression of sFlt1 led to pronounced reductions in CVP forming endothelial cells (**J**). (**L**, **M**) Quantification of the number of endothelial cells that comprise the CVP at 72 (**L**) and 144 (**M**) hpf. (**N**–**Q**) 6 dpf WT (**N**), *vegfab^-/-^;vegfd^-/-^* (**O**), *vegfab^-/-^;vegfc^-/-^* (**P**), and *vegfab^-/-^;vegfd^-/-^;vegfc^-/-^* (**Q**) CVP visualized by *Tg(kdrl:*EGFP) expression after immunostaining for GFP. (**R**) Quantification of the number of endothelial cells that comprise the CVP at 6 dpf (the number of animals examined per genotype is listed in the panel). (**S**) Schematic representations of the CVP phenotypes observed in *vegfa* mutants or after sFlt1 overexpression. Zebrafish *vegfa* paralog mutants genetically interact in fenestrated CVP formation. Temporal inhibition of Vegfa signaling by sFlt1 overexpression recapitulated the severely impaired CVP phenotypes observed in genetic mutants, providing evidence for well conserved mechanisms of fenestrated CNS vessel development between zebrafish and mammals Scale bars: 50 μm in **D** for **A**–**D**, in **K** for **H**–**K**, in **Q** for **N**–**Q**.

To inhibit Vegfa signaling in a temporally-controlled manner, we employed the previously established *Tg(hsp70l:sflt1)* line in which the Vegfa ligand trap, sFlt1, is overexpressed upon a heatshock treatment (*37*, *38*). Since active angiogenesis of choriocapillaris formation occurs between 36 and 65 hpf in zebrafish embryos (*36*), we subjected embryos to heatshocks at 34, 44, and 54 hpf, and analyzed them at 72 hpf (Fig. 6f). Similar to what we observed in *vegfab* and *vegfaa* mutants, we found a drastic reduction in the number of endothelial cells that comprise the choriocapillaris after sFlt1 overexpression (Fig. 6H, 6J). This phenotype was not observed in sibling controls, or *Tg(hsp70l:sflt4)* animals in which the Vegfc/d ligand trap, sFlt4, overexpression was induced (*37*, *38*), under the same time course of heatshock treatments (Fig. 6I, 6K). To eliminate the possibility of potential developmental delays in heatshock treated *Tg(hsp70l:sflt1)* larvae, we also analyzed them at 144 hpf, following additional heatshocks every 24 hours after 54 hpf (Fig. 6G). The results were similar even at this later stage (Fig. 6M), demonstrating that inhibition of Vegfa-induced signaling at the time of choriocapillaris formation is sufficient to abrogate this process.

The results of heatshock treated *Tg(hsp70l:sflt4)* larvae indicate that Vegfc and/or Vegfd themselves do not contribute to choriocapillaris development. Nevertheless, we analyzed these mutants individually or in combinations to determine their definite contributions to choriocapillaris formation. We observed that *vegfc* and *vegfd* single mutants or *vegfc;vegfd* double mutants showed either no significant difference or a slight reduction in the number of endothelial cells that comprise the choriocapillaris at 6 dpf (Fig. 6R). However, we found no genetic interactions between *vegfab* and *vegfd* or *vegfc* in choriocapillaris formation (Fig. 6N–R). This finding, combined with the lack of significant changes in endothelial cell numbers of the choriocapillaris following sFlt4 overexpression, suggest that Vegfc or Vegfd contributions to choriocapillaris development are minor, if any.

Together, our observations here reveal that the combined deletion of *vegfab, vegfc*, and *vegfd* does not increase the severity of defects in fenestrated choriocapillaris formation compared to *vegfab* single mutants, in contrast to our recent finding in mCP vascularization. Instead, *vegfab* and *vegfaa* display a strong genetic interaction in choriocapillaris formation, which is further supported by the results from sFlt1-mediated temporal Vegfa inhibition experiments (Fig. 6S). Combined with *vegfab* and *vegfaa* restricted expression in RPE, these data suggest highly conserved cellular and molecular mechanisms of choriocapillaris development between zebrafish and mammals (*33*, *34*). Moreover, these results provide further evidence for regional heterogeneity of angiogenic regulation across fenestrated CNS vascular beds since the deletion of the same ligand combination surprisingly caused no fenestrated vessel formation defect at the dCP/PG interface (Fig. 3N).

### Endothelial cell-type-specific angiogenic programs drive fenestrated capillary development at the hypophysis and OVLT

The hypophysis (pituitary gland) is the vital neuroendocrine organ consisting of two lobes, the adenohypophysis (anterior pituitary) and the neurohypophysis (posterior pituitary) (*39*). The neurohypophysis (NH) forms a neurovascular interface where neuropeptides, such as oxytocin and vasopressin, are secreted into blood circulation via fenestrated capillaries, which regulate reproduction, fluid balance, and blood pressure (*39*–*41*). As we presented earlier (Fig. 1P–W), zebrafish larvae deficient for Wnt/β-catenin signaling components such as Gpr124, Reck, and Wnt7aa, do not exhibit an apparent defect in fenestrated capillary formation at the hypophysis and around the OVLT, a sensory CVO critical for body fluid homeostasis via osmotic sensing and regulation (*42*–*44*). Prior work implies that the OVLT resides dorsally to the anterior part of hypophyseal artery (HyA) (*45*). To date, angiogenic cues that drive fenestrated capillary development in these CVOs remain unclear.

A previous study characterized the developmental process of capillary loop formation at the hypophysis (*46*). HyA formation begins by sprouting from the cranial division of the internal carotid artery toward the midline, which extends rostorally until it connects with the palatocerebral arteries (PLA) at around 60 hpf (*46*). On the other hand, hypophyseal veins (HyV) start sprouting medially from the bilateral primary lateral vein sinuses at around 60 hpf and position in the posterior part of the vascular loop at the hypophysis (Hy loop). Bilateral HyA form the anterior portion of the loop and fuse with HyV to develop the Hy loop by 72 hpf (*46*). Ultrastructural analysis of the pituitary gland in both larval and adult zebrafish revealed vascular endothelial fenestrations in this organ (*47*, *48*). In support of this observation, we noted strong *Tg(plvap:EGFP)* fenestrated marker expression in endothelial cells that comprise the Hy loop and HyA at 120 hpf (Fig. 1P), although much fainter *Tg(plvap:EGFP)* expression was seen in HyV.

We examined expression patterns of *vegfaa, vegfab, vegfc*, and *vegfd* using BAC transgenic reporters at several developmental stages and observed distinct and partially overlapping expression patterns in the ventral diencephalon. Prominent and specific *vegfaa* expression was detected in cells that reside in close proximity to the Hy loop (Fig. 7A, 7E). Strong *vegfab* expression was found in cells that lie rostrodorsally to the anterior part of the Hy loop (Fig. 7B, 7F) and also in cells that reside at the junction where HyA extends to join the PLA (Fig. 7G). Notable *vegfc* expression was observed in several different cell types that reside closely to the anterior portion of the HyA (Fig. 7C, 7H). Expression of *vegfd* BAC reporter was not detectable in this brain region at the developmental stages we examined (Fig. 7D). Moreover, *vegfaa, vegfab*, and *vegfc* BAC reporter expression was not observed in endothelial cells that constitute HyA, HyV, or Hy loop.

**Figure 7.**
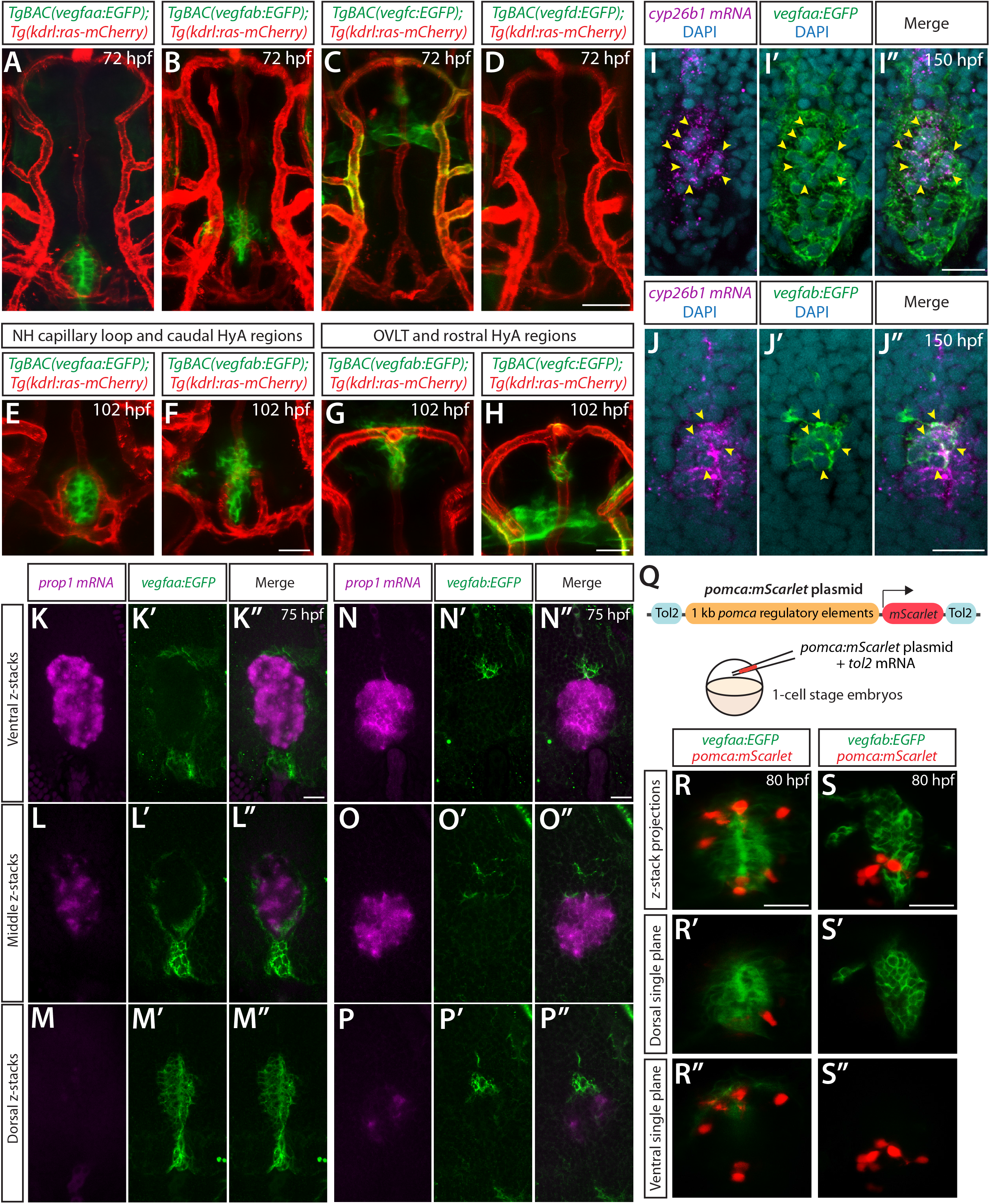
BAC transgenic analysis of *vegf* expression at the developing hypophysis and OVLT. (**A**–**D**) Dorsal views of 72 hpf *TgBAC(vegfaa:EGFP)* (**A**), *TgBAC(vegfab:EGFP)* (**B**), *TgBAC(vegfc:EGFP)* (**C**), and *TgBAC(vegfd:EGFP)* (**D**) ventral brain vasculature of larvae that carried the *Tg(kdrl:ras-mCherry)* endothelial transgene. Prominent *TgBAC(vegfaa:EGFP)* and *TgBAC(vegfab:EGFP)* expression was observed in cells that reside in close proximity to capillary loop at the neurohypophysis (**NH**) (**A**, **B**). Notable *TgBAC(vegfc:EGFP)* expression was detected in the anterior portion of the hypophyseal artery (**HyA**) around the organum vasculosum of the lamina terminalis (**OVLT**) (**C**). *TgBAC(vegfd:EGFP)* expression was not detectable at this stage (**D**). (**E**–**H**) Magnified images of 102 hpf *TgBAC(vegfaa:EGFP)* (**E**), *TgBAC(vegfab:EGFP)* (**F**, **G**), and *TgBAC(vegfc:EGFP)* (**H**) ventral brain vasculature of larvae that carried the *Tg(kdrl:ras-mCherry)* transgene. *TgBAC(vegfaa:EGFP)^+^* or *TgBAC(vegfab:EGFP)^+^* cells lie slightly dorsal to NH capillary loop (**E**, **F**) with many *TgBAC(vegfab:*EGFP)^+^ cells being localized further rostrodorsally likely in the hypothalamus. In rostral HyA regions around the HyA-PLA junction and OVLT, peri-vascular *TgBAC(vegfab:EGFP)* and *TgBAC(vegfc:EGFP)* expression was detected (**G**, **H**). Additionally, strong *TgBAC(vegfc:EGFP)* signals were observed around the OVLT. (**I**–**I”**) Single confocal z-plane images of 150 hpf *TgBAC(vegfaa:EGFP)* larval ventral brain following *in situ* hybridization of *cyp26b1*, showing overlapping signals between the EGFP^+^ cells and *cyp26b1^+^* pituicyte (n=10). Yellow arrowheads indicate the overlapping cells. (**J**–**J”**) Single confocal z-plane images of 150 hpf *TgBAC(vegfab:EGFP)* larval ventral brain following *in situ* hybridization of *cyp26b1*, showing overlapping signals between the EGFP^+^ cells and *cyp26b1^+^* pituicyte (n=11). Yellow arrowheads indicate the overlapping cells. (**K**–**M”**) Serial confocal z-stacks of ventral (**K**–**K”**), middle (**L**–**L”**), and dorsal (**M**–**M”**) images showing no overlapping signals between *TgBAC(vegfaa:EGFP)^+^* and *prop1^+^* cells by *in situ* hybridization in 75 hpf larvae (n=8). (**N**–**P”**) Serial confocal z-stacks of ventral (**N**–**N”**), middle (**O**–**O”**), and dorsal (**P**–**P”**) images showing no overlapping signals between *TgBAC(vegfab:EGFP)^+^* and *prop1^+^* cells by *in situ* hybridization in 75 hpf larvae (n=10). (**Q**) Schematic of the *pomca:mScarlet* construct used for injection experiments (**R**–**S”**). (**R**–**S”**) Magnified dorsal views of 80 hpf *TgBAC(vegfaa:EGFP)* (**R**–**R”**) and *TgBAC(vegfab:EGFP)* (**S**–**S”**) NH of larvae that were injected with the *pomca:mScarlet* construct at the one-cell stage. Confocal z-stack maximum projection (**R**, **S**), dorsal (**R’**, **S’**), and ventral (**R”**, **S”**) single z-plane images showing no overlapping signals between EGFP^+^ cells and *pomca:*mScarlet^*+*^ pituitary corticotrophs. Scale bars: 50 μm in **D** for **A**–**D**; 25 μm in **F** for **E**–**F**, in **H** for **G**–**H**, in **R** for **R**–**R”**, in **S** for **S**–**S”**; and 15 μm in **I”** for **I**–**I”**, in **J”** for **J**–**J”**, in **K”** for **K**–**M”**, in **N”** for **N**–**P”**.

To determine cell type(s) that express these *vegf* genes, we carried out the following experiments. First, we performed *in situ* hybridization on these BAC transgenic reporters using cell type-specific markers. We used a *cyp26b1* probe, which marks the astroglial pituicyte, and a *prop1* probe for progenitors, as used in recent studies (*47, 49, 50)*. Although we did not observe any overlap between *prop1^+^* progenitors and *vegfaa* or *vegfab* BAC reporter^+^ cells at the hypophysis (Fig. 7K–P”), we found significant co-localizations between *cyp26b1^+^* pituicytes and both of these reporter^+^ cells (Fig. 7I–J” and Figure 7–figure supplement 1), suggesting that pituicytes are a source of these Vegf ligands. These results are consistent with the recently published bulk and single-cell RNA-seq data that show *vegfaa* and *vegfab* expression in pituicytes (*47, 49, 50)*. Additionally, we injected a plasmid construct in which mScarlet expression is driven under the *pomc* promoter that was previously characterized to induce gene expression in pituitary corticotrophs (*51*). We observed no co-localization between *pomc^+^* cells and *vegfaa^+^* or *vegfab^+^* cells at the hypophysis (Fig. 7O–S”), indicating that pituitary corticotrophs are not a cell type that expresses these two *vegf* genes. A series of these co-localization experiments suggest that pituicytes are the major cell type expressing *vegfaa* and *vegfab* at the hypophysis. Importantly, scRNA-seq data of the adult neurohypophysis in mice showed pituicyte-specific expression of *Vegfa*, indicating interspecies conservation of *vegfa* expression in pituicytes (*52*).

Next, we analyzed *vegfaa, vegfab, vegfc*, and *vegfd* mutants individually or in various combinations to determine the requirements of Vegf ligands for pituitary gland vascularization. We noted that none of these individual *vegf* mutants displayed an obvious defect in the formation of the HyA, Hy loop, or HyV (Fig. 8A–D, 8I). Although the combined deletion of *vegfab, vegfc*, and *vegfd* caused no defect in the HyV and Hy loop, we identified some triple mutants (7 out of 18, approximately 39%) exhibiting either absence or partial formation of the HyA (Fig. 8H, 8I). A comparable phenotype was observed even in *vegfab;vegfc* double mutants (7 out of 22, approximately 32%) (Fig. 8G, 8I), suggesting that this HyA formation defect results from a genetic interaction between *vegfab* and *vegfc*. Contribution of Vegfd to this vascularization process is likely absent or minor since no genetic interaction between *vegfd* and *vegfab* or *vegfc* was noted (Fig. 8E, 8F, 8I). These genetic data are in line with undetectable expression of *vegfd* BAC reporter.

**Figure 8.**
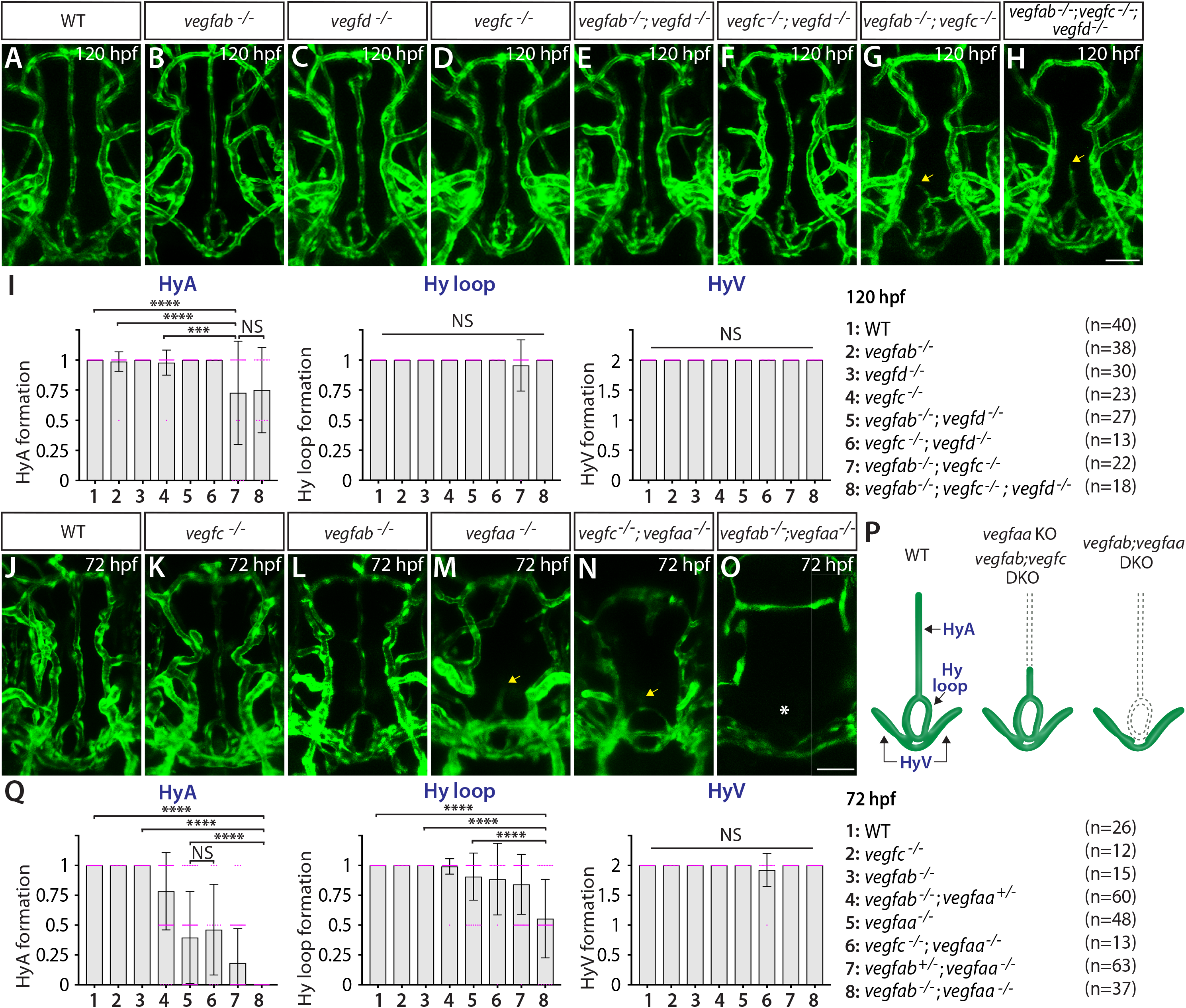
Local regulatory heterogeneity of Vegfs-dependent angiogenesis at the neurohypophysis and OVLT. (**A**–**H**) Dorsal views of 120 hpf WT (**A**), *vegfab^-/-^* (**B**), *vegfd^-/-^* (**C**), *vegfc^-/-^* (**D**), *vegfab^-/-^;vegfd^-/-^* (**E**), *vegfc^-/-^;vegfd^-/-^* (**F**), *vegfab^-/-^;vegfc^-/-^* (**G**), and *vegfab^-/-^;vegfc^-/-^;vegfd^-/-^* (**H**) ventral brain vasculature visualized by *Tg(kdrl*:EGFP) expression. A significant fraction of *vegfab^-/-^;vegfc^-/-^* (**G**) and *vegfab^-/-^;vegfc^-/-^;vegfd^-/-^* (**H**) larvae exhibited a partial formation of the HyA, resulting in HyA stalling at the halfway point towards the PLA (arrows, **G**, **H**). (**I**) Quantification of HyA, Hy loop, and HyV formation at 120 hpf (the number of animals examined per genotype is listed in the panel). *vegfab^-/-^;vegfc^-/-^* and *vegfab^-/-^;vegfc^-/-^;vegfd^-/-^* larvae displayed a specific and partially penetrant defect in HyA formation. (**J**–**O**) Dorsal views of 72 hpf WT (**J**), *vegfc^-/-^* (**K**), *vegfab^-/-^* (**L**), *vegfaa^-/-^* (**M**), *vegfc^-/-^;vegfaa^-/-^* (**N**), and *vegfab ^-/-^vegfaa^-/-^* (**O**) ventral brain vasculature visualized by *Tg(kdrl:EGFP)* expression. Similar to *vegfab^-/-^;vegfc^-/-^* larvae, *vegfaa^-/-^* fish exhibited a specific and partially penetrant defect in HyA formation, leading to HyA stalling at the halfway point towards the PLA (arrow, **M**). The severity of this phenotype was exacerbated by the simultaneous deletion of *vegfab*, but not *vegfc* (arrow, **N**), showing a genetic interaction between *vegfaa* and *vegfab*, but not between *vegfaa* and *vegfc*. Intriguingly, phenotypes in *vegfab^-/-^;vegfaa^-/-^* larvae were restricted to the fenestrated HyA and Hy loop (asterisk, **O**) with no significant defect in HyV formation. (**Q**) Quantification of HyA, Hy loop, and HyV formation at 72 hpf. Scale bars: 50 μm in **H** for **A**–**H** and in **O** for **J**–**O**.

Since *vegfab;vegfc;vegfd* triple or *vegfab;vegfc* double mutants displayed only a partially penetrant defect in HyA formation, we next investigated the role of Vegfaa in this process. Interestingly, we found that *vegfaa* single mutants exhibited either absent or stalled HyA phenotypes (38 out of 48, approximately 79%) (Fig. 8M), similar to *vegfab^-/-^;vegfc^-/-^* larvae, but to a greater extent. However, this HyA formation defect in *vegfaa* single mutants is still partially penetrant (Fig. 8Q), allowing us to test combined deletions of *vegfaa, vegfab*, and/or *vegfc ^hu6410^*. We observed that the severity of HyA stalling phenotype in *vegfaa^-/-^* larvae was substantially exacerbated by the simultaneous deletion of *vegfab*, but not of *vegfc*. These results show that there is a genetic interaction between *vegfaa* and *vegfab*, but not between *vegfaa* and *vegfc* (Fig. 8N–Q). Intriguingly, we noted that the phenotypes observed in *vegfab^-/-^;vegfaa^-/-^* larvae were restricted to fenestrated Hy loop and the HyA with no defect in HyV formation (Fig. 8O, 8Q). This finding is in stark contrast to the dCP/PG interface where BBB-forming MsV was selectively impaired without any noticeable defect in fenestrated vessel development in *vegfab^-/-^;vegfaa^-/-^* larvae (Fig. 3N). The selective loss of fenestrated Hy loop and the HyA in *vegfab^-/-^;vegfaa^-/-^* larvae suggest that phenotypically distinct Hy loop/HyA and HyV depend on discrete angiogenic mechanisms (Fig. 8P).

Lastly, we tested whether temporal inhibition of Vegfa signaling could recapitulate this fenestrated angiogenesis defect observed in *vegfaa;vegfab* double mutants. To this aim, we employed the *Tg(hsp70l:sflt1)* and *Tg(hsp70l:sflt4)* lines, heatshocked embryos at 34, 44, and 54 hpf, and analyzed larvae at 72 or 144 hpf (Fig. 9A). We observed that heatshock-induced overexpression of sFlt1 led to stalled HyA formation at the halfway point towards the PLA, accompanied by a drastic reduction in the number of endothelial cells that comprise the HyA (Fig. 9B–D, 9F, 9G). Milder, yet robust, reduction of endothelial cell numbers in the Hy loop was detected in heatshock-treated *Tg(hsp70l:sflt1)* larvae, but only smaller differences were recognized in the HyV of these larvae. In contrast, sFlt4 overexpression led to no difference in the number of endothelial cells that comprise the Hy loop and HyV compared to controls, while we observed a significant reduction in HyA endothelial cell numbers at 144 hpf (Fig. 9E–G). These sFlt4 overexpression results indicate Vegfc’s selective contribution to endothelial cell proliferation in HyA, which is in line with the specific defect in HyA formation observed in *vegfab;vegfc* double mutants. Thus, temporal inhibition of Vegfa or Vegfc/d signaling during vascularization of the hypophysis/OVLT is sufficient to cause HyA and Hy loop formation defects similarly to what was observed in the mutants.

**Figure 9.**
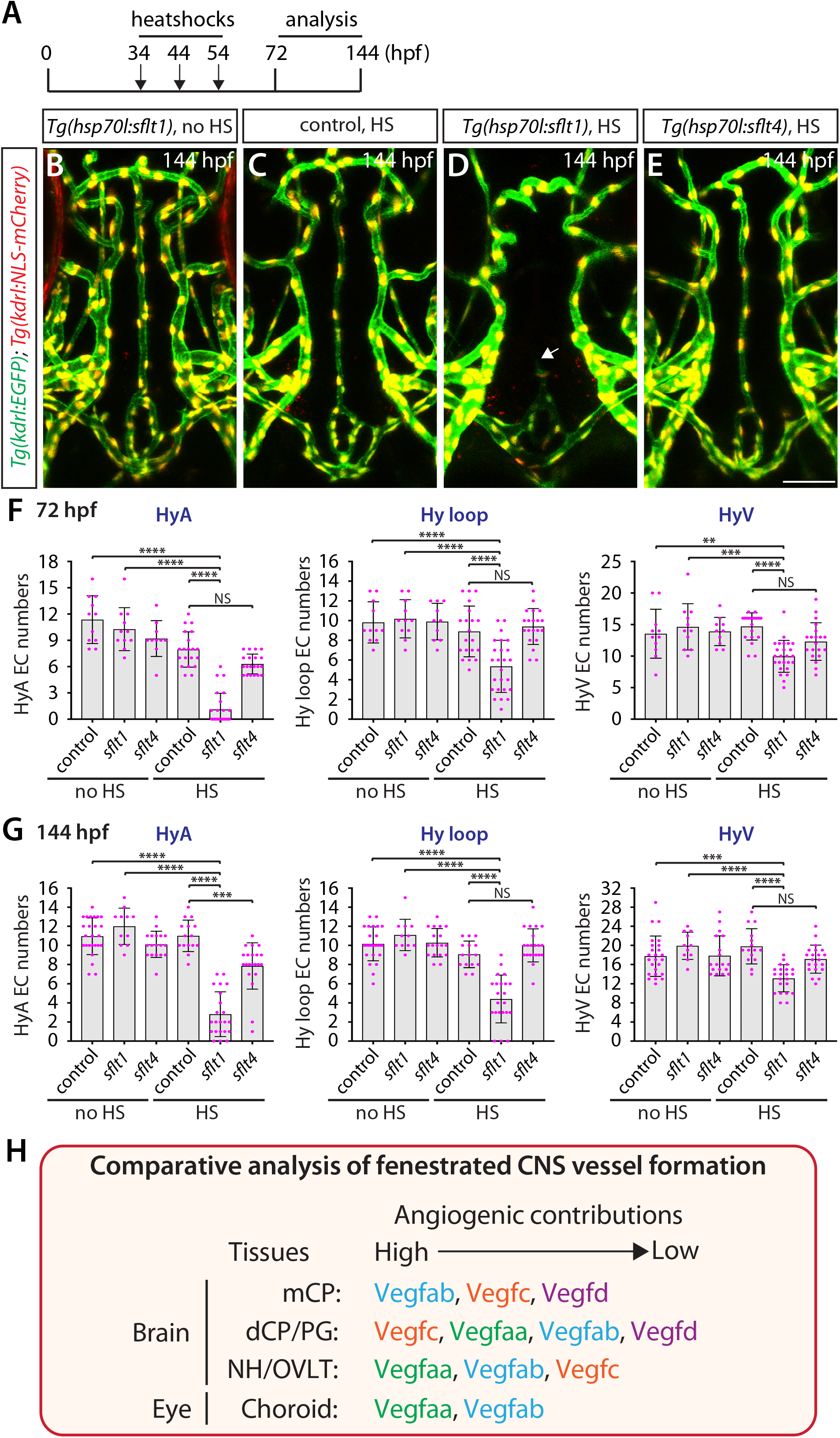
Temporal inhibition of Vegfa signaling by sFlt1 overexpression is sufficient to cause stalled HyA and impaired Hy loop phenotypes at the neurohypophysis and OVLT. (**A**) Experimental time course of the heat-shock treatments for panels (**B**–**G**). *Tg(hsp70l:sflt1), Tg(hsp70l:sflt4)*, and their control siblings were subjected to a heat-shock at 34, 44, and 54 hpf, and analysed 72 or 144 hpf. (**B**–**E**) Dorsal views of 144 hpf *Tg(hsp70l:sflt1);Tg(kdrl:EGFP);Tg(kdrl:NLS-mCherry)* (**B**, **D**), *Tg(kdrl:EGFP);Tg(kdrl:NLS-mCherry)* (**C**), and *Tg(hsp70l:sflt4);Tg(kdrl:EGFP);Tg(kdrl:NLS-mCherry)* (**E**) ventral brain vasculature after treatment with (**C**–**E**) and without (**B**) multiple heatshocks. Heatshock-induced overexpression of sFlt1 caused severe defects in HyA formation, leading to HyA stalling at the halfway point towards the PLA (arrow, **D**). (**F**, **G**) Quantification of the number of endothelial cells that comprise the HyA, Hy loop, and HyV at 72 (**F**) and 144 (**G**) hpf. Heatshock-induced overexpression of sFlt1 led to a drastic reduction in the number of endothelial cells that comprise the HyA. Milder, yet robust, reduction of endothelial cell numbers in Hy loop was detected in heatshock-treated *Tg(hsp70l:sflt1)* larvae, but only smaller differences were recognized in the HyV of these larvae. In contrast, sFlt4 overexpression led to no difference in the number of endothelial cells that comprise the Hy loop and HyV compared to controls. However, we observed a significant reduction in HyA endothelial cell numbers at 144 hpf (**G**), indicating Vegfc’s selective contribution to endothelial cell proliferation in the HyA. (**H**) Summary table depicting the results from our comparative analysis of fenestrated CNS vessel formation in this and previous studies. Angiogenic contributions were speculated based on the severity and penetrance of observed phenotypes and also on the levels of genetic interactions detected to cause phenotypes. This comparative data uncovers regional heterogeneity of Vegfs-mediated angiogenic regulation and reveals an unexpected, crucial role for Vegfc in fenestrated brain vascular development. Scale bar: 50 μm in **E** for **B**–**E**.

Collectively, our results here present vessel type-selective vascularization mechanisms. Vegfa paralogs and Vegfc selectively regulate the formation of fenestrated Hy loop and the HyA with little contribution to HyV development. These observations highlight local regulatory heterogeneity of Vegf-mediated angiogenesis and requirements of diverse genetic programs needed for vascularization across fenestrated CNS vascular beds.

## DISCUSSION

Fenestrated and BBB-forming capillaries mediate brain-blood communications crucial for diverse neural activities and brain homeostasis. Despite the recent great advances in BBB brain angiogenesis, fenestrated capillary development in the CPs and CVOs has been poorly understood. Here, we report substantial functional redundancy among Vegf ligands and their molecular identity required for fenestrated capillary development in multiple CNS vascular beds. We show that the individual loss of four different Vegf ligands (Vegfaa, Vegfab, Vegfc, and Vegfd) in zebrafish causes either undetectable or partially penetrant defects in fenestrated vessel formation in the CPs, PG, hypophysis, OVLT, and retinal choroidal vascular plexuses. However, the simultaneous loss of specific Vegf ligand combinations results in dramatically enhanced phenotypic defects (Fig. 9H). Expression analysis using BAC transgenic reporters supports the genetic mutant results by uncovering partially overlapping expression of relevant *vegf* genes at the time of vessel morphogenesis. In each of the developing fenestrated CNS vascular beds we examined, relevant *vegf* gene expression is observed in extending endothelial cells and/or non-neuronal cell types that are uniquely present in the CPs, CVOs, and retina. These non-neuronal cell types include epithelial cells in the CPs, RPE in the outer retina, and astroglial pituicytes in the NH. Genetic analysis of the mutant allele *vegfc^um18^*, which lacks Vegfc paracrine activity, provides evidence for endothelial cell-autonomous and non-autonomous contributions of Vegfc angiogenic activity to fenestrated vessel formation at the CP/PG interface. Heatshock-induced overexpression of sFlt1, a decoy receptor for Vegfa ligands, implies that RPE-derived Vegfaa and Vegfab play a crucial role in fenestrated choriocapillaris formation. Altogether, our comparative results presented here provide valuable insights into a core angiogenic mechanism underlying fenestrated CNS vascular development.

### Capillary type-selective angiogenesis directed by Vegfs and Wnt7/β-catenin signaling

How phenotypically-heterogeneous networks of brain vasculature arise remains unclear. We recently reported that a redundant angiogenic activity of Vegfab, Vegfc, and Vegfd is restricted to fenestrated mCP vascular development with little impact on the formation of the neighboring BBB vasculature (*12*). In contrast, we presented here that the genetic loss of Wnt7/β-catenin signaling (*gpr124, reck*, or *wnt7aa* KO) causes a severe angiogenesis defect in the brain parenchyma without any apparent deficit in fenestrated vasculature across the CNS. This observation is in line with the recent reports showing that β-catenin activation is maintained at remarkably low levels in fenestrated vascular beds of the CPs and CVOs compared to those that form the BBB (*53*, *54*). Our further observations of selective defects in fenestrated PrA and HyA formation in *vegfab^-/-^;vegfc^-/-^* larvae at the dCP/PG interface and hypophysis, respectively, provide additional evidence for endothelial cell-type-selective angiogenic deficits. Moreover, a selective loss of the BBB-forming vessel MsV in *vegfab^-/-^;vegfaa^-/-^* larvae without a defect in the neighboring fenestrated vasculature at the dCP/PG interface is another striking example of endothelial cell-type-specific angiogenesis. Notably, *vegfab^-/-^;vegfaa^-/-^* larvae display pronounced defects that are restricted to fenestrated Hy loop and the HyA in the NH, presenting heterogeneous requirements of the same Vegf ligand combination in BBB and fenestrated angiogenesis across organs.

Together, these findings support a model whereby endothelial fates are pre-determined prior to angiogenesis, leading to differential or heterogeneous responses of adjacent endothelial cells to angiogenic cues presented in local microenvironments. This differential endothelial responsiveness, in turn, results in the specification of distinct fates and properties. Intriguingly, these observations imply that brain endothelial cell fates are imprinted before they migrate to their target tissues, presenting a model for the fate determination of brain endothelium and the induction of vessel type-selective angiogenesis.

### Local and inter-tissue heterogeneity of Vegfs-mediated angiogenic regulation during the vascularization of fenestrated CNS vascular beds

Current evidence suggests that the development of complex brain vascular networks requires extensive functional redundancy among signaling components/pathways (*55*). For example, brain vasculature that forms the BBB is not uniformly compromised in mice deficient for Wnt/β-catenin signaling (*22*, *24*, *55*). This is because Wnt ligands (Wnt7a, Wnt7b, Norrin) and their receptors display significant redundancy in BBB formation and maintenance across distinct regions of the brain (*18*, *22*, *24*, *55*–*57*). Wnt7a and Wnt7b are redundantly required for the vascularization of the ventral neural tube in mice (*22*, *24*). Wnt7a/b and Norrin genetically interact in regulating BBB maintenance in the cerebellum (*55*). These prior studies have indicated brain region-specific molecular requirements for vascularization and barrier properties, which can involve multiple redundant molecular cues.

In this study, we began by asking whether the combination of Vegf ligands (Vegfab, Vegfc, and Vegfd) required for complete vascularization of the mCP is the master regulator of fenestrated capillary development across the brain. To our surprise, our comparative analysis of the CPs and CVOs reveals remarkable heterogeneity of angiogenic requirements within and across these organs. This local and inter-tissue heterogeneity of Vegfs-mediated angiogenic regulation during the vascularization of fenestrated brain vascular beds is supported by brain area-specific, unique gene expression of these angiogenic factors. In all the brain regions we examined, we observed partially overlapping and distinct *vegf* expression patterns in either developing endothelial cells or non-neuronal cell types that specifically reside in/around the CPs or CVOs.

How are fenestrated angiogenic programs encoded in a brain region-specific manner? In the zebrafish NH, specialized astroglia, pituicytes, were implicated as a potential source of Vegfab and Vegfaa, which maintain high vascular permeability in this region (*47*), consistent with our observations of these two genes’ expression in pituicytes at early larval stages. In the CPs of adult mice, epithelial cells express high levels of VegfA (*58*) that maintain the fenestrated states of vasculature in this region (*58*, *59*). Our present study suggests that epithelial cells express *vegfab* and *vegfc* in the developing dCP, which together direct the vascularization of the CP/PG interface. Given these observations, we speculate that non-neuronal cell types present in the CPs and CVOs build unique endothelial features to meet specific needs in these brain regions. Additionally, it is a possibility that brain region-specific angiogenic modulators are present to shape spatiotemporal activities of redundant angiogenic cues.

### Identification of Vegfc/d as crucial regulators of fenestrated brain vessel formation

An unexpected finding made through our comparative analysis is that fenestrated vessel formation requires Vegfc/d across multiple brain regions. In contrast to the well-characterized function of VEGFA in angiogenesis, endothelial fenestration, and vascular permeability both *in vitro* and *in vivo* (*60*–*62*), a regulatory role for Vegfc/d in this context is largely unclear *in vivo*. Furthermore, a role for Vegfc/d in brain vascularization has not been well established. It is known that *vegfc* loss-of-function leads to a transient, partially penetrant defect in the formation of the primordial hindbrain channel in early embryonic stages of zebrafish (*31*, *63*) and also that Vegfc/d contribute to mCP vascularization (*12*). Additionally, previous work documented the expression of VEGFC and its major receptor VEGFR3 in multiple neuroendocrine organs and their fenestrated capillaries in human tissues (*64*). Consistent with this report, our *vegfc* BAC transgenic zebrafish reporter marks developing fenestrated capillaries in addition to cells that reside within several neuroendocrine organs, including the pineal and pituitary gland. These observations imply a conserved function of Vegfc in regulating fenestrated capillary formation/integrity across vertebrates.

Our findings raise many questions that remain unanswered. We noted that *vegfc* displays a strong genetic interaction with *vegfa* paralog(s) to drive fenestrated vessel formation across the CPs and CVOs. Does Vegfc play a crucial role in endothelial fenestration development and/or maintenance? Does the redundancy between Vegfc and Vegfa paralog(s) occur at the level of their receptor(s) and downstream signaling pathways, or through a genetic compensatory mechanism? Is proteolytic activation of Vegfc involved in fenestrated brain vessel development? If so, what are these proteolytic regulators and where are they expressed? Is Vegfc function in this context conserved in mammals? Addressing these questions will further advance our understanding of Vegfc function in brain vascularization and barrier properties.

In summary, defining unique sets of molecular cues required for the formation and maintenance of specialized vascular beds within the brain is fundamental to designing vascular bed-specific cerebrovascular therapies. Given a wide range of organs that form fenestrated vasculature outside of the CNS, including in endocrine glands (pancreas and thyroid), kidney, and intestine, our findings may have broad impacts on our understanding of fenestrated endothelial development and specializations beyond the CNS. Finally, to extend our work in zebrafish, future investigation of the Vegfc/d signaling axis as a potential modulator of fenestrated vessel formation in mammals is warranted.

## Abbreviations

ACeV: Anterior cerebral vein
BBB: Blood-brain barrier
CNS: Central nervous system
CPs: Choroid plexuses
CrDI: Cranial division of the internal carotid artery
CVOs: Circumventricular organs
CVP: Choroidal vascular plexus
Cyp26: Cytochrome P450 family 26 enzymes
DLV: Dorsal longitudinal vein
Glut1b: Glucose transporter 1b
HyA: Hypophyseal artery
Hy loop: Hypophyseal loop
HyV: Hypophyseal veins
MCeV: Middle cerebral vein
MsV: Mesencephalic vein
NH: Neurohypophysis
OA: Optic artery
OVLT: Organum vasculosum of the lamina terminalis
PCeV: Posterior cerebral vein
PG: Pineal gland
PGV: Pineal gland vessel
PLA: Palatocerebral arteries
PrA: Prosencephalic artery
Prop1: Paired-Like Homeobox 1
Plvap: Plasmalemma vesicle-associated protein
Pomca: Proopiomelanocortin a
vECs: Vascular endothelial cells
Vegfs: Vascular endothelial growth factors

## Acknowledgements

We thank Drs. Didier Stainier, Nathan Lawson, and Michael Taylor for kindly providing us with fish lines; Don Zeisloft and his team for zebrafish care and husbandry; Michelle America and Leslie Sanderson for assistance of zebrafish mutant maintenance; Dr. Judith Drazba and her team for confocal and electron microscopy imaging. This work was supported by funding from the NIH (R01 NS117510) and start-up funds from the Cleveland Clinic Foundation to R.L.M.

## Author contributions

S.P. and R.L.M. designed experiments and wrote the manuscript; S.P., O.A.C., Q.C., L.D.B., R.E.Q., W.H., and R.L.M. performed experiments and analyzed data. B.V. generated and provided critical zebrafish lines for conducting this study. G.L. provided unpublished bioinformatics information and supported experiments. All authors commented on the manuscript.

## Data and materials availability

Correspondence and requests for materials should be addressed to Ryota L. Matsuoka.

## Experimental Procedures

### Zebrafish husbandry and strains

All zebrafish husbandry was performed under standard conditions in accordance with institutional and national ethical and animal welfare guidelines. All zebrafish work was approved by the Cleveland Clinic’s Institutional Animal Care and Use Committee under the protocol number 00002684. The following lines were used in this study: *Tg(kdrl:EGFP)^s843^* (*65*); *Tg(kdrl:Has.HRAS-mcherry)^s896^* (*66*), abbreviated *Tg(kdrl:ras-mCherry); Tg(kdrl:NLS-mCherry)^is4^* (*67*); *Tg(UAS:EGFP-CAAX)^m1230^* (*68*); *Tg(UAS-E1b:NfsB-mCherry)^c264^* (*69*), abbreviated *Tg(UAS:NTR-mCherry); Tg(hsp70l:sflt1, cryaa-cerulean)^bns80^* (*37*), abbreviated *Tg(hsp70l:sflt1); Tg(hsp70l:sflt4, cryaa-cerulean)^bns82^* (*37*), abbreviated *Tg(hsp70l:sflt4)*; *TgBAC(vegfab:gal4ff)^bns273^* (*27*); *TgBAC(vegfaa:gal4ff)^lri96^* (*12*); *TgBAC(vegfc:gal4ff)^bns270^* (*12*); *TgBAC(vegfd:gal4ff)^lri95^* (*12*); *Tg(glut1b:mCherry)^sj1^* (*70*); *Tg(plvapb:EGFP)^sj3^* (*70*), abbreviated *Tg(plvap:EGFP); Et(cp:EGFP)^sj2^* (*17*); *vegfaa^bns1^* (*28*); *vegfab^bns92^* (*28*); *vegfc^hu6410^* (*71*); *vegfc^um18^* (*31*); *vegfd^bns257^* (*72*); *gpr124 ^s984^* (*19*); and *wnt7aa ^ulb2^* (*20*). Adult fish were maintained on a standard 14 h light/10 h dark daily cycle. Fish embryos/larvae were raised at 28.5°C. To prevent skin pigmentation, 0.003% phenylthiourea (PTU) was used beginning at 10-12 hpf for imaging. Fish larvae analyzed at 10 dpf were transferred to a tank containing approximately 250 mL water supplemented with 0.003% PTU (up to 25 larvae/tank) and fed with Larval AP100 (<50 microns dry diet, Zeigler) starting at 5 dpf.

### Genotyping of mutants

Genotyping of *vegfaa^bns1^* and *vegfab^bns92^* mutant fish was performed by high-resolution melt analysis of PCR products as described previously(*12*). Genotyping of *vegfc^hu6410^*, *vegfc^um18^*, and *vegfd^bns257^* mutant fish was performed by standard PCR as described previously(*12*, *72*). Genotyping of *gpr124 ^s984^* and *wnt7aa ^ulb2^* mutant fish was performed by high-resolution melt analysis of PCR products using the following primers:

*gpr124 s984* forward: 5’ – AGGGTCCACTGGAACTGCACACATTG – 3’

*gpr124 s984* reverse 5’ – TGCAATGGAAAGGCAGCCTGTCTCC – 3’

*wnt7aa ulb2* forward: 5’ – GCGCAAATGGGAATCAATGAGTG – 3’

*wnt7aa ulb2* reverse: 5’ – TCCAAAGACAGTTCTTTCCCCGAG – 3’

### Generation and genotyping of *reck* mutants

The zebrafish *reck^ulb3^* allele was engineered by targeted genome editing using the CRISPR/Cas9 system, as described previously(*20*, *73*). sgRNA target site was designed using the CRISPR Design website (http://crispr.mit.edu). The oligos (5-taggCCTGACAGTACTCACGAC -3’ and 5’-aaacGTCGTGAGTACTGTCAGG -3’) were annealed and ligated into the pT7-gRNA vector (Addgene #46759)(*74*) after digesting the plasmid with BsmBI (NEB). The sgRNA was transcribed from the BamHI linearized pT7-gRNA vector using the MEGAshortscript T7 kit (Thermo Fisher Scientific). The synthetic Cas9 mRNA was transcribed from the XbaI linearized pT3TS-nls-zCas9-nls vector (Addgene #46757)(*74*) using the mMESSAGE mMACHINE T3 Kit (Ambion). sgRNA (30 pg) and nls-zCas9-nls mRNA (150 pg) were injected into one-cell stage zebrafish embryos. The *reck^ulb3^* mutant allele harbors a 12 base pair deletion in the exon that encodes part of the 3rd cysteine knot motif. Please refer to Figure 1–-figure supplement 1 for detailed information. Genotyping of *reck^ulb3^* mutant fish was performed by high-resolution melt analysis of PCR products using the following primers:

*reck ulb3* forward: 5’ – TTACGCAGGCAGACACACCACCTG – 3’

*reck ulb3* reverse: 5’ – GAGACGGTGGGAGACGAGTCTGTG – 3’

### High-resolution melt analysis (HRMA)

A CFX96 Touch Real-Time PCR Detection System (Bio-Rad) was used for the PCR reactions and high-resolution melt analysis. Precision melt supermix for high-resolution melt analysis (Bio-Rad) was used in these experiments. PCR reaction protocols were: 95°C for 2 min, 46 cycles of 95°C for 10 s, and 60°C for 30 s. Following the PCR, a high-resolution melt curve was generated by collecting EvaGreen fluorescence data in the 65–95°C range. The analyses were performed on normalized derivative plots.

### Heat shock treatments

Fish embryos/larvae raised at 28.5°C were subjected to a heat shock at 37°C for 1h by replacing egg water with pre-warmed (37°C) egg water, as described previously(*37*, *38*). After each heat shock, the fish embryos/larvae were kept at room temperature for 10 min to cool down and then incubated at 28.5°C. Heat shock experiments were performed as follows: *Tg(hsp70l:sflt1, cryaa-cerulean)^bns80^, Tg(hsp70l:sflt4, cryaa-cerulean)^bns82^*, and their sibling animals without the *cryaa:*cerulean eye marker, which carried the reporters *Tg(kdrl:EGFP)^s843^;Tg(kdrl:NLS-mCherry)^is4^*, were subjected to a heat shock at 34, 44, and 54 hpf, and imaged at 72 or 144 hpf, for vascular analysis in the hypophysis and OVLT brain regions. For analysis of choriocapillaris formation, the same Tg lines were employed and subjected to either a heat shock at 34, 44, and 54 hpf and imaged at 72 hpf or a heatshock at 34, 44, 54, 78, 102, 126 hpf and imaged at 144 hpf.

### Immunohistochemistry

Immunohistochemistry was performed by following standard immunostaining procedures as described previously (*12*). For CVP analysis of 72 hpf or 6 dpf larvae, fish larvae were fixed in 4% paraformaldehyde (PFA)/phosphate buffered saline (PBS) overnight at 4°C and dehydrated through immersion in methanol serial dilutions (50%, 75%, then 100% methanol three times, 10 min each) at room temperature (RT). The dehydrated samples were stored in 100% methanol at −20°C before eye dissections. Microdissections of only one eye per fish was carried out under SMZ18 stereomicroscope (Nikon) at 5-8X magnification using Dumont forceps. Dissected eyes were collected for immunostaining while the remaining tissues were collected for genotyping when necessary. The following primary antibodies were used: chicken anti-GFP (Aves Labs at 1:1000), rabbit anti-DsRed (Clontech at 1:300), mouse anti-Claudin-5 (4C3C2, Invitrogen at 1:500), and mouse anti-ZO-1 (ZO1-1A12, Thermo Fisher Scientific at 1:500).

### *in situ* hybridization combined with immunostaining in zebrafish larvae

Whole-mount mRNA *in situ* hybridization was conducted by following the protocol as previously described (*75*). Briefly, digoxigenin (DIG) labelled RNA probes were achieved by reverse transcription and purification of the amplified partial coding sequence of selected genes with a DIG RNA labelling kit (Roche, Switzerland). For 75 and 150 hpf larvae, 30- and 40-min proteinase K digestion was conducted, respectively. Probes for chain hybridization reaction (HCR) *in situ* hybridization v3.0 were ordered from Molecular Instruments (USA). The protocol of HCR *in situ* hybridization on whole-mount zebrafish larvae was slightly modified from the previously described protocol (*76*, *77*). 8 pmol probes were used for each gene and samples were fixed in 4% PFA/PBS and washed between detection and amplification steps. Immunostaining with anti-eGFP (GFP Rabbit Polyclonal #A11122 or GFP Chicken Polyclonal #A10262, Invitrogen, Thermo Fisher Scientific, USA) was probed after *in situ* hybridization as described previously (*52*). The larvae were then mounted in 75% glycerol and imaged with a Zeiss LSM980 confocal microscope. Single plane and serial z-stack images were extracted using Fiji-ImageJ software (Fiji, Japan).

### Plasmid injections

For *pomca:mScarlet* plasmid injection, the plasmid was generated by cloning the mScarlet coding sequence downstream of 1,006 bp *pomca* regulatory elements (*47*, *51*) flanked by Tol2 sites. The 1,006 bp *pomca* regulatory elements were amplified by PCR from genomic DNA of zebrafish embryos using the following primers:

*pomca* promoter forward: 5’- atta<u>aagctt</u>CTTTTTTATTCTGCTTTAAGACCTC -3’

*pomca* promoter reverse: 5’- atta<u>gaattc</u>CTCTGAGAACACAATACAATTCAC -3’

Hind III and EcoR I restriction enzyme sites (underlined) were incorporated into the primers to allow cloning. The engineered construct (30 pg per embryo) was injected together with transposase mRNA (*tol2* mRNA, 25 pg per embryo) into *Tg(UAS:EGFP-CAAX)^m1230^* one-cell stage embryos that carry the *TgBAC(vegfab:gal4ff)^bns273^* or *TgBAC(vegfaa:gal4ff)^lri96^* transgene.

### Confocal and stereo microscopy

A Leica TCS SP8 confocal laser scanning microscope (Leica) was used for live, immunofluorescence, and transmitted differential interference contrast imaging. Fish embryos and larvae were anaesthetized with a low dose of tricaine, embedded in a layer of 1% low melt agarose in a glass-bottom Petri dish (MatTek), and imaged using 10X dry or 25X water immersion objective lens. Leica Application Suite X (LAS X) software (Version 3.7.0.20979) was used for image acquisition and analysis.

### Transmission electron microscopy

10 dpf wild-type larvae were anesthetized with tricaine and fixed by immersion in 2.5% glutaraldehyde/4% paraformaldehyde/0.2 M sodium cacodylate (pH 7.4) for 2 days at 4°C. Samples were post-fixed in 1% osmium tetroxide for 1 h and incubated in 1% uranyl acetate in Maleate buffer for 1 h. Samples were dehydrated through immersion in methanol serial dilutions and embedded in epoxy resin. Ultrathin sections of 85 nm were cut with a diamond knife, collected on copper grids, and stained with uranyl acetate and lead citrate. Images were captured using a transmission electron microscope (FEI Tecnai G2 Spirit BioTWIN).

### Quantification of DLV, PCeV, MCeV, and MsV formation

Fish carrying the *Tg(kdrl:EGFP)^s843^* reporter were used for this quantification. Fish larvae at indicated developmental stages were analyzed for the presence or absence of the DLV. To quantify bilaterally formed PCeV, MCeV, and MsV, the following criteria were used to score the extent of each vessel’s formation: 1) Score 2 - when the bilateral vessels are both fully formed; 2) Score 1.5 - when the vessel on one side is fully formed, but that on the other side is partially formed; 3) Score 1 - when the vessel on one side is fully formed, but that on the other side is absent; 4) Score 0.5 - when the vessel on one side is partially formed, but that on the other side is absent; and 5) Score 0 - when the bilateral vessels are both absent.

### Quantification of PrA, PGV, and ACeV at the dCP/PG interface

Fish carrying the *Tg(kdrl:EGFP)^s843^* reporter were used for this quantification. Fish larvae were analyzed at 72 hpf or 10 dpf for the extent of vascular formation at the dCP/PG interface. To quantify bilaterally formed PrA, PGV, and ACeV, the same scoring criteria used to quantify the extent of bilateral vessel formation (PCeV, MsV, and MCeV) described earlier were applied.

### Quantification of endothelial cells that comprise the CVP

Fish carrying *Tg(kdrl:EGFP)^s843^; Tg(kdrl:NLS-mCherry)^is4^* or *Tg(kdrl:EGFP)^s843^* reporter were used to quantify the number of endothelial cells that formed the CVP. Fish larvae were fixed at 72 hpf or 6 dpf, dehydrated through methanol serial dilutions, subjected to eye dissections, and then immunostained for GFP, DsRed, and ZO-1, a marker for RPE. Immunostained eyes were imaged from the back of the eyes to capture the CVP that resides outside of the RPE layer. *Tg(kdrl:EGFP)^+^* or *Tg(kdrl:*NLS-mCherry)^+^ cells that resided outside of the RPE layer and that lied directly adjacent to RPE were defined as CVP endothelial cells and counted.

### Quantification of HyA, Hy loop, and HyV

Fish carrying the *Tg(kdrl:EGFP)^s843^* reporter were used for this quantification. Fish larvae were analyzed at 72 or 120 hpf for the extent of hypophyseal vascular formation. To quantify bilaterally formed HyV, the same scoring criteria used to quantify the extent of bilateral vessel formation (PCeV, MsV, and MCeV) described earlier were applied.s HyA formation was quantified as follows: 1) score 1 -when the entire vessel is fully formed with vessel extension from the Hy loop to the PLA; 2) score 0.5 - when the vessel formation is partial, failing to reach the PLA; and 3) score 0 - complete absence of the vessel. The Hy loop was quantified as follows: 1) score 1 - when the entire capillary loop is fully formed; 2) score 0.5 - when capillary loop formation is partial, lacking a part of the capillary loop; and 3) score 0 - when the loop is completely absent.

### Statistical analysis

Statistical differences for mean values among multiple groups were determined using a one-way analysis of variance (ANOVA) followed by Tukey’s multiple comparison test. Fisher’s exact test was used to determine significance when comparing the degree of penetrance of observed phenotypes. The criterion for statistical significance was set at *P* < 0.05. Error bars are SD.

---

**Supplemental Movie 1. Fenestrated vasculature at the PG/dCP interface.**

10 dpf *Et(cp:EGFP);Tg(kdrl:ras-mCherry)* head immunostained for rhodopsin shows the 3D spatial relationship between dCP epithelial cells (green), pineal photoreceptor cells (magenta), and blood vessels (white).

**Figure 1– figure supplement 1.**
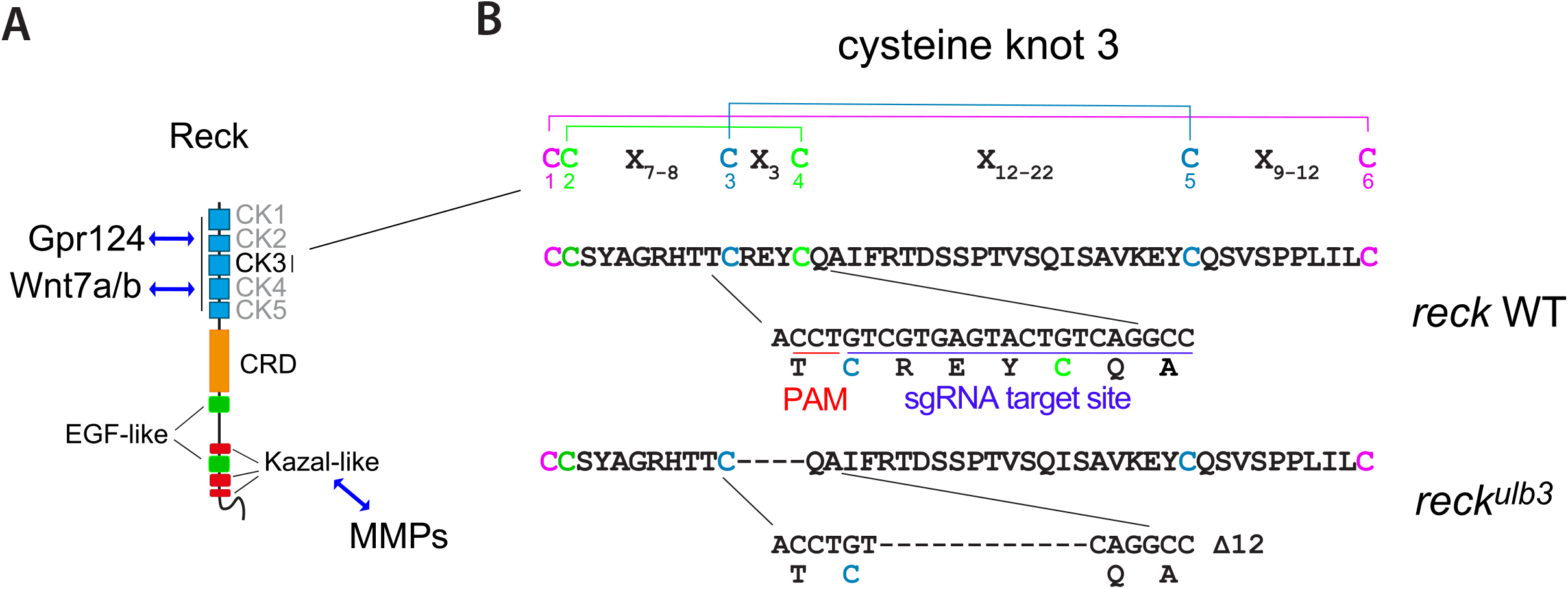
Characterization of the *reck^ulb3^* allele. (**A**) Schematic representation of the domain architecture of Reck, with from N- to C-terminus, five cysteine knot motifs (CK), a Frizzled-like cysteine-rich domain (CRD), two Epidermal Growth Factor-like domains (EGF-like) and three Kazal-like motifs upstream of a membrane glycosylphosphatidylinositol (GPI) anchor. The CK motifs are implicated in Wnt signaling by binding Gpr124 and Wnt7a/b, while the Kazal motifs control matrix metalloproteinase (MMP) activity. (**B**) Sequence alignment of WT and mutant Reck sequences showing the disulfide bonding pattern within the cysteine knot motif consensus (6 Cys repeat: C-C-X_7-9_-C-X_3_-C-X_12-22_-C-X_9-12_-C). The *reck^ulb3^* mutant allele was generated by CRISPR-Cas9 mutagenesis at the target site highlighted in blue, see Methods. PAM: protospacer adjacent motif. An in-frame mutation allele was selected to interfere selectively with the domains of Reck implicated in Wnt-dependent brain angiogenesis.

**Figure 7– figure supplement 1.**
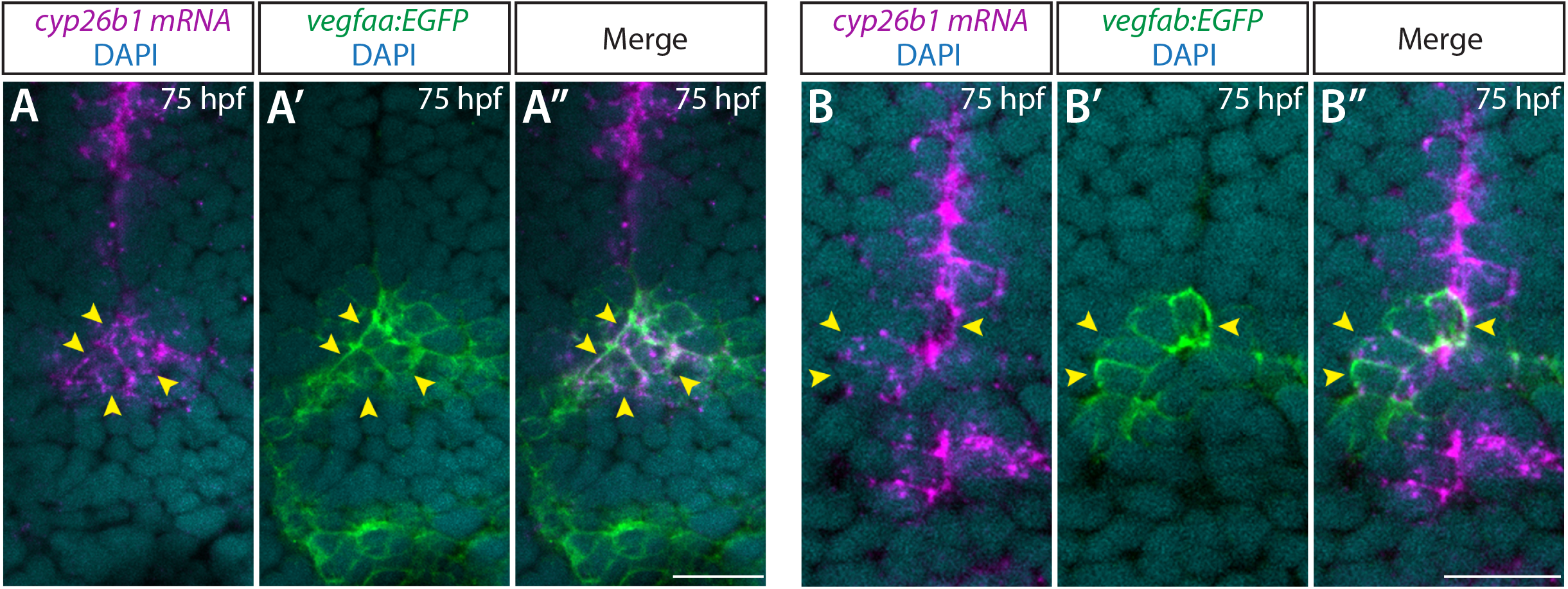
Co-localization of the pituicyte marker *cyp26b1* and *vegfaa* or *vegfab* BAC transgenic reporter expression at early larval stages. (**A**–**A”**) Single confocal z-plane images of 75 hpf *TgBAC(vegfaa:EGFP)* larval ventral brain following *in situ* hybridization of *cyp26b1*, showing overlapping signals between the EGFP^+^ cells and *cyp26b1^+^* pituicyte (n=10). Yellow arrowheads indicate the overlapping cells. (**B**–**B”**) Single confocal z-plane images of 75 hpf *TgBAC(vegfab:EGFP)* larval ventral brain following *in situ* hybridization of *cyp26b1*, showing overlapping signals between the EGFP^+^ cells and *cyp26b1^+^* pituicyte (n=11). Yellow arrowheads indicate the overlapping cells.

